# Microbial symbionts are shared between ants and their associated beetles

**DOI:** 10.1101/2022.12.02.518891

**Authors:** Catalina Valdivia, Justin A. Newton, Sean O’Donnell, Christoph von Beeren, Daniel J. C. Kronauer, Jacob A. Russell, Piotr Łukasik

**Author notes:** Correspondence, Catalina Valdivia, Symbiosis Evolution Research Group, Institute of Environmental Sciences, Jagiellonian University ul. Gronostajowa 7, 30-387 Kraków, Poland.

## Abstract

The transmission of microbial symbionts across individuals and generations can be critical for animal development and survival. Likewise, the transmission of microbes across closely interacting species could also affect host biology. Army ants (Formicidae: Dorylinae) and their hundreds of closely associated insect species (myrmecophiles) can provide a unique insight into interspecific symbiont sharing. Here, we compared the microbiota of workers and larvae of the army ant *Eciton burchellii* with those of 13 myrmecophile beetle species using 16S rRNA amplicon sequencing. We found that the previously characterized symbionts of army ant workers (Unclassified Firmicutes and Unclassified Entomoplasmatales) were largely absent from ant larvae and from myrmecophiles, whose microbial communities were usually dominated by *Rickettsia, Wolbachia, Rickettsiella*, and/or *Weissella*. Strikingly, different species of myrmecophiles and ant larvae often shared identical 16S rRNA genotypes of common bacteria. In particular, army ant larvae, some workers, and several myrmecophile species often hosted identical *Weissella* (Lactobacillales), based on 16S rRNA and also protein-coding gene sequences. Also, we found high relatedness between some newly characterized *Weissella* and animal-associated strains from aquatic and marine habitats. Looking more broadly, we found *Weissella* OTUs in 11.6% of samples from nearly all habitats and environments characterized by the Earth Microbiome Project. Together, our data show that unrelated but closely interacting species can share much of their microbiota. The high relatedness of strains found across such disparate hosts as ants, beetles, trout, and whales suggests that some versatile microbes move between hosts and habitats despite few opportunities for direct interaction.

## Introduction

Host-associated microbiota, or the communities of microbes occupying defined habitats in host bodies, have played important roles in animal ecology and evolution (McFall-Ngai et al., 2013). Symbiotic microbes have enabled the emergence and evolutionary success of multiple host clades and species, and strongly affected individual host fitness and population dynamics (Fisher et al., 2017). The effects of symbiotic microbial strains on hosts range from positive nutritive or defensive interactions to harmful parasitic or pathogenic relationships (Berg et al., 2020). These effects vary across host and symbiont genotypes and environmental conditions (Sze et al., 2020). There are relatively clear differences among the functional categories of symbiotic microbes: closed, mixed and open (Perreau & Moran, 2021). Closed symbioses are characterized by the presence of microbes transmitted strictly vertically (maternally) across host generations. They include obligate intracellular symbionts of insects that feed on nutritionally incomplete diets such as plant sap or blood, and require supplementation with essential amino acids and vitamins (Bennett & Moran, 2015; Rio, Attardo, & Weiss, 2016). Microbes that form “mixed” symbioses, in addition to utilizing vertical transmission, can also transmit horizontally, i.e., move among non-relatives within and across host species. These associations include facultative endosymbiotic bacteria of insects such as *Wolbachia* (Kaur et al., 2021), which can range in their effects from beneficial to deleterious, depending on conditions. In the third major category, open symbiosis, exemplified by most gut microbiomes, microbes are generally transmitted through the environment. While primarily linked to nutrition, gut microbiota also play other critical roles, such as protection against parasites or pathogens, or influencing development (Engel & Moran, 2013).

The factors shaping the effects of microbial symbioses across different host species are not well understood, with research strongly biased towards vertebrate and especially mammalian systems and their “open” symbioses (Petersen & Osvatic, 2018). However, in taxa such as insects, it may be more common to find microbes that establish mixed or closed relationships. How the microbial symbionts are transmitted across individuals of a species largely determines opportunities for their interspecific transfer. In mammals, the acquisition of microbes needed for development or survival mostly occurs during birth and early life, through shared environment (Campos-Cerda & Bohannan, 2020; Ferretti et al., 2018; Moeller et al., 2018). In insects, mechanisms for symbiont maternal transmission, including transovarial transmission, the deposition of symbiont-containing capsules, egg smearing with fecal matter (Ohbayashi et al., 2020), or through manipulating nest environment (Shukla et al., 2018), have evolved repeatedly. On the other hand, social behavior is also important, particularly for the transmission of gut microbiota. Through shared environment, shared food sources, or feeding habits (e.g. coprophagy or trophallaxis), microbial communities of social animals more closely resemble those of the other members of their own social group (colony, family, herd, etc.) than those of other groups (Bo et al., 2020; Brito et al., 2019; Engel & Moran, 2013; Sarkar et al., 2020).

Sharing the same environment by different species may provide opportunities for interspecific transmission of gut microbes. The similarities in microbial community composition between humans and their pet dogs (Song et al., 2013) serve as a good example. Likewise, there are reports of microbe sharing across interacting insect species. For example, the similarities in gut microbiota composition between fungus-growing ants and their social parasites in another ant genus were explained by nest space and food sharing, or myrmecophile predatory behavior (Liberti et al., 2015), and it has been shown that velvety tree ants and their mymercophiles have similarities in microbiota composition (Perry et al., 2021). On the other hand, many insects lack abundant gut microbiota, with facultative endosymbionts often dominating the microbial communities (Hammer et al., 2019). The horizontal transmission of these mixed symbionts may be more challenging, as they are often unable to survive outside of the host environment; at the same time, due to their often high abundance in insect tissues, they may be easier to study. There are reports of apparent transmission of *Wolbachia* from *Drosophila simulans* to the parasitic wasp *Leptopilina boulardi* (Heath et al., 1999), from prey to predator in the case of mites and terrestrial isopods (Clec’h et al., 2013; Hoy & Jeyaprakash, 2005), and from ants to kleptoparasitic ant crickets (Orthoptera: Myrmecophilidae - Tseng et al., 2020) - even if the phylogenetic resolution of the presented data did not always suffice to confirm the patterns. Overall, it appears that the likelihood of interspecific transmission should correlate with the frequency and intensity of the interactions among host insect species.

Army ants (Formicidae: Dorylinae) in the genus *Eciton* are a particularly interesting group from the social interaction perspective (Kronauer, 2020). With each colony numbering thousands to millions of workers, these nomadic ants roam terrestrial habitats in search of animal prey. They practice dependent colony founding: new colonies form by fission when a large colony splits in two, each retaining a single queen (Gotwald, 1995; Peeters & Ito, 2001; Schneirla, 1971). These colonies host a diverse set of closely associated invertebrate species, collectively known as myrmecophiles (Akre & Rettenmeyer, 1968). Myrmecophilous insects, which include beetles, especially in the family Staphylinidae (rove beetles; (Parker, 2016; Von Beeren et al., 2021), vary in their level of integration and roles in army ant colonies (Akre & Rettenmeyer, 1968; Rettenmeyer, 1961; von Beeren et al., 2018; von Beeren, Maruyama et al., 2011). Some of these insects, whether living within colonies or staying on their outskirts, eat the food the ants collect or the waste they produce, acting as commensals (Rettenmeyer, 1961). Others prey on larvae or worker ants of the host colony, while still others use the ants merely as a means of transport (Gotwald, 1995; Rettenmeyer et al., 2011; Rettenmeyer, 1961). These diverse interactions create multiple opportunities for the transmission of microbes among community members: either directly between species, or through their shared social environment. However, despite the ecological importance of army ants as keystone species in tropical forests (Hoenle et al., 2019; Kronauer, 2020; Pérez-Espona et al., 2018), we know relatively little about the dynamics, specificity, or sharing of their microbial symbionts. It was shown that the microbiota of New World army ant workers consist primarily of two specialized bacteria, Unclassified Firmicutes and Unclassified Entomoplasmatales (Anderson et al., 2012; Funaro et al., 2011; Łukasik et al., 2017; Mendoza-Guido et al., 2022, in press), but little is known about the microbial associations of other community members, including army ant larvae and myrmecophiles.

The goal of this project was to assess the degree of microbial overlap across unrelated but cohabiting species inside *Eciton burchellii parvispinum* (Winston et al., 2017) army ant colonies. We addressed it by sequencing 16S rRNA gene amplicons for 105 beetles representing 13 species, from eight different colonies from a Costa Rican montane forest site, in addition to ant workers and larvae. We then tracked microbial clades and genotypes across species and colonies. For the most broadly distributed microbial taxon, the genus *Weissella* (Lactobacillales), we increased the resolution of strain association analysis by sequencing a protein-coding gene. Finally, after realizing that insect-associated *Weissella* strains can closely resemble vertebrate-associated strains from marine/aquatic habitats, we looked at the distribution of *Weissella* in different environments using data from the largest global collection of diverse samples published so far, the Earth Microbiome Project (Thompson et al., 2017). By combining these data, we show substantial overlap in microbial composition among different hosts and a broad environmental distribution of *Weissella*.

## Material and methods

### Insect collection and identification

We sampled insects in July 2012 in the rainforest of Monteverde, Costa Rica (Fig. 1A, Table S2). From raiding or emigration columns of eight *Eciton burchellii parvispinum* colonies, we collected medium-sized ant workers, myrmecophilous beetles and, in two cases, army ant larvae. We also captured six presumedly free-living staphylinid beetles in leaf-litter away from the sampled army ant colonies. Upon collection, specimens were immediately preserved in 95% ethanol and stored at −20°C until processing. There was one exception: nine myrmecophiles from two morpho-species were starved for 24 hours prior to preservation, in order to test the persistence of microbes within myrmecophiles.

**Figure 1.**
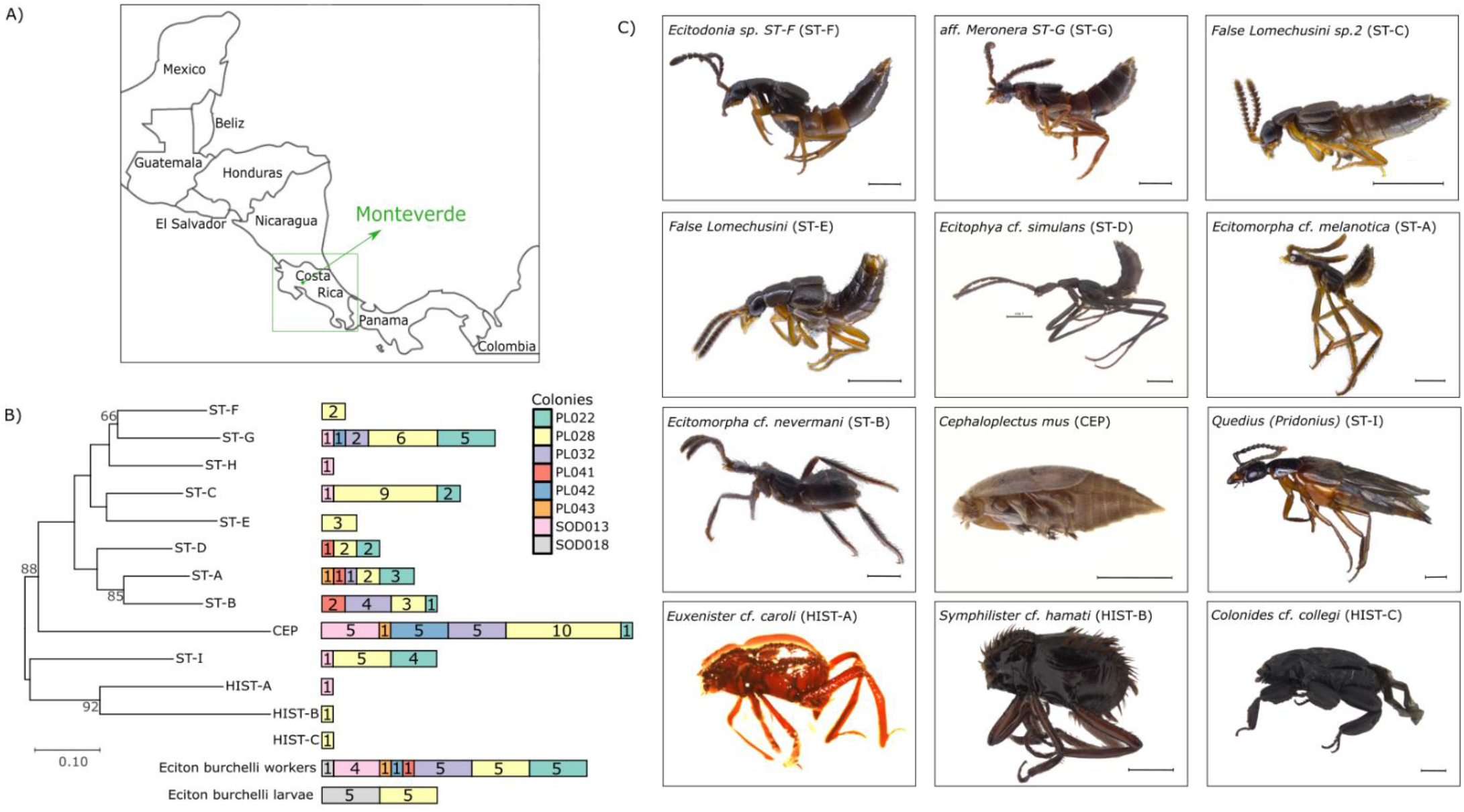
The origin and diversity of the studied myrmecophiles. A) The sampling location;B) A maximum-likelihood tree for twelve myrmecophilous beetle species used in this study, based on a 658bp portion of the mitochondrial cytochrome oxidase I (*COI*) gene. Bars represent the number of specimens representing thirteen species, or ant workers/larvae, obtained from different E. burchellii colonies. In the tree, bootstrap support values >60% are shown. C) Representative specimens of twelve myrmecophile species. Scale bars represent 1 mm.

We had previously identified all ant colonies based on morphological characters of workers (Longino, 2010) and the partial sequence of the mitochondrial cytochrome oxidase I (*COI*) gene (Łukasik et al., 2017). The same methods were used to categorize myrmecophilous beetles. In short, specimens were individually mounted and z-stack images were produced using a Leica Z16 APO stereomicroscope equipped with a light dome, a Leica DFC450 camera, and the processing software Leica application suite (version 4) at The Rockefeller University. This allowed a preliminary identification to morphospecies based on morphological features (Fig. 1C). Within a few hours, specimens were placed back into ethanol and stored at −20°C, limiting DNA degradation or microbial colonization. For a subset of these specimens, we then sequenced the *COI* gene, as explained below. Using the latest species identification keys for each morphospecies and a recently published *COI* reference database of *Eciton* myrmecophiles (Von Beeren et al., 2021), we identified all myrmecophiles to the lowest taxonomic level possible. In cases where we were not able to identify the species, we added our morphotype designation to the taxon name to provide unique identifiers (e.g., *Ecitodonia sp. ST-F*). Details on species identifications of myrmecophiles are provided in Supplementary Appendix. The species distribution across the sampled colonies is shown in Fig. 1B.

Some specimens had *COI* signals that did not match their morphospecies but instead matched with possible prey or parasite sequences from the NCBI database. We made a note of this to check if these *COI* signals had some influence on the results from the 16S rRNA sequence analysis.

### Symbiont screens and amplicon library preparation

Prior to DNA extraction, all insects were surface-sterilized through 1-minute immersion in 1% bleach, followed by rinsing with molecular-grade water. We extracted DNA from dissected gasters (ant workers) or whole specimens (larvae, beetles) using DNeasy Blood and Tissue kits (Qiagen Ltd.), following the protocol for Gram-positive bacteria. The DNA extractions were used for PCR reactions with the universal primers 9Fa and 907R (Łukasik et al., 2017; Russell et al., 2009) for the bacterial 16S rRNA gene and LCO-1490 and HCO-2198 (Folmer et al., 1994) for the *COI* gene of ants and myrmecophiles. PCR reaction conditions were described previously (Lukasik et al 2017). The 16S rRNA gene amplification success, assumed to correlate with the bacterial abundance in the sample, was estimated by comparing the brightness of bands in an agarose gel against negative extraction controls. If the brightness of 16S rRNA bands was equal to or lower than that of these negative controls, such samples were not processed further.

DNA samples that were classified as having substantial bacterial load were submitted, in three separate batches, including also other sample types, to Argonne National Laboratory for the preparation of amplicon libraries and subsequent sequencing of the V4 hypervariable region of the 16S rRNA gene (primers: 515F-806R), following the Earth Microbiome Project protocols (Caporaso et al., 2012). Paired-end, 150bp sequencing was performed on an Illumina MiSeq platform. Along with insect DNA samples, six extraction blanks and three blanks consisting of molecular grade water were sequenced to aid in identifying contaminants.

### Microbiota data analysis

The amplicon data were analyzed using a custom protocol combining vsearch and usearch with custom Python scripts, described in detail at https://github.com/catesval/army_ant_myrmecophiles. Briefly, after extracting reads corresponding to the experimental libraries from the sequencing runs and adding data for previously characterized ant specimens (Łukasik et al., 2017), we quality-filtered and assembled reads into contigs using PEAR v0.9.11 (Zhang et al., 2014). We performed dereplication using vsearch v2.15.2_linux_x86_64 (Rognes et al., 2016) and denoising with usearch v11.0.667_i86linux32 (Edgar, 2010; Edgar et al., 2011). After processing all samples, separately in order to avoid the loss of rare genotypes during denoising, zOTU lists were merged. We then performed OTU picking and chimera removal using algorithms incorporated in the usearch software. Next, taxonomy was assigned using vsearch, using the SILVA SSU 138 database as reference. Finally, we used custom scripts to merge the outputs of OTU picking and taxonomy assignment to create the OTU and zOTU tables.

As shown previously, contamination during sample processing can strongly alter the microbial community profiles of organisms with less abundant microbiota, including some army ants (Łukasik et al., 2017; Salter et al., 2014). Because of this, we screened and filtered putative contaminant genotypes, identified based on their relative abundance in libraries representing insect samples and negative controls. We adapted and expanded Łukasik at al. 2017’s custom script that considers three criteria. First, our script compared the maximum relative abundance of unique genotypes in libraries representing insect samples against the maximum relative abundance of these same genotypes in blank libraries. If the ratio of these values was below the specified threshold (here, set to 10), then the genotype was classified as a contaminant and removed. Second, the script classified any genotype that reached a relative abundance threshold (here: 0.001) as “symbiont”; we concluded that the remaining microbial signal, likely a combination of rare symbiotic microbes, uncommon contaminants, and sequencing artifacts, cannot be classified reliably, but included them as “others” in relative abundance and other comparisons. Finally, the script classified libraries as “heavily contaminated” and removed them from the list if the proportion of reads classified as contaminants exceeded a third threshold (here: 0.8). While this approach should effectively eliminate contaminants, its downside is the possible exclusion of rare symbiotic microbes.

However, in our analyses, we focused on abundant and widespread microbes. Specifically, we selected for more detailed investigation those 97% OTUs that fulfilled at least two of the following three criteria: the average relative abundance of the OTU across samples was equal to or higher than 0.01; its relative abundance in at least one sample was ≥ 0.05; or it was present in at least 20 samples.

For the calculation of the genotype-level composition (zOTU) of the OTUs, we calculated what percentage of each 97% OTU was represented by different zOTUs, and then filtered zOTUs to keep only those that were represented in the dataset by at least 100 reads and present with an abundance of at least 0.05 in at least one sample. The final data, managed using Microsoft Excel, were visualized using the pheatmap library (Kolde, 2019) in R version 3.6.3 (2020-02-29).

### *rplB* genotype-level diversity of *Weissella*

To obtain additional insights into the diversity of *Weissella*, one of the most broadly distributed symbionts in our dataset, we amplified and sequenced a portion of the *50S ribosomal protein L2* (*rplB*) gene from 51 Monteverde specimens in which the microbe was present (Figure 2, Table S4). Additionally, 31 samples from other army ant species and collection sites that were not a part of the primary dataset but tested positive for *Weissella* based on specific PCR primers or amplicon data (Łukasik et al. 2017), were included for *rplB* sequencing. To achieve this, we used newly designed primers within the *rplB* and the adjacent *rpsS* gene: ArWei_rplB_F3: GGTCGTCGTAATATGACTGGT, Leu_rpsS_R3: TGAACGACGTGACCATGTCTTG, and Wei_RpsS_Rseq: CTTCAACCTTCTTCTTCAACAACAAGYKRGC. The PCR program was: 94°C for 1 minute, 25 cycles of 95<u>°</u>C for 15 seconds, 70<u>°</u>C->62.8C (decreasing by 0.3<u>°</u>C each cycle) for 15 seconds, 72C for 20 seconds; 35 cycles of 94C for 15 seconds, 58C for 15 seconds, 72<u>°</u>C for 20 seconds; 70<u>°</u>C for 2 minutes. The *rplB* sequences obtained after trimming rpsS fragments and intergenic region were aligned against publicly available homologs identified in the NCBI database via BLASTn (Table S6).

**Figure 2.**
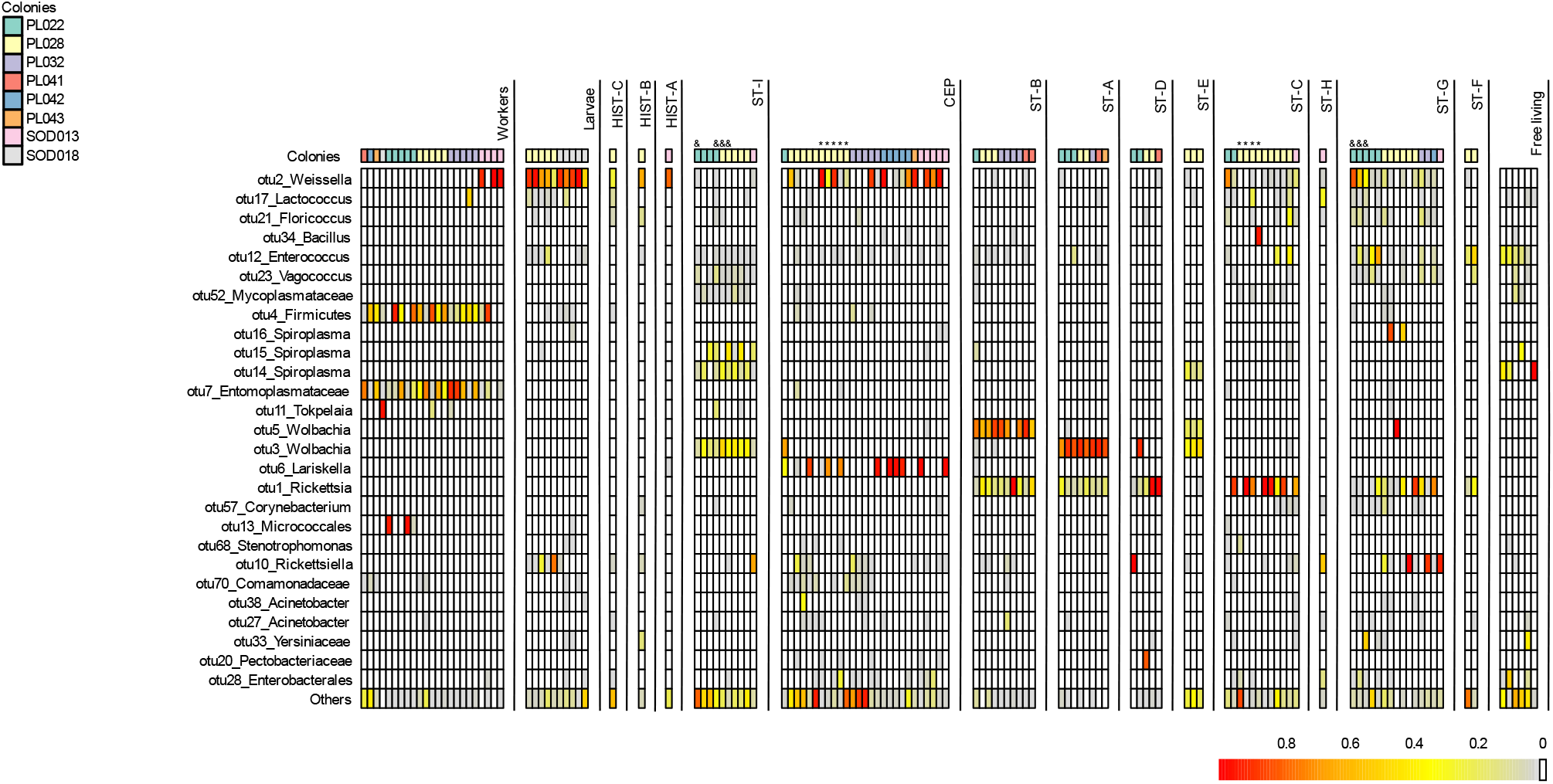
The relative abundance of the microbial 97% OTUs in specimens of *E. burchellii* workers, larvae, and 13 myrmecophile species from eight ant colonies from Monteverde, and a few free-living staphylinids. Columns represent insect specimens divided into species, ordered based on *COI* ML tree, and labeled on top with the abbreviation assigned in Fig.1. Rows represent bacterial OTUs sorted according to ML phylogeny for their representative genotypes, with bootstrap values over 60 shown. The color gradient represents the relative abundance of each OTU (rows) in each of the insect specimens (columns). All OTUs with relative abundance values.

### Phylogenetic analyses

We quality-checked and aligned all insect *COI* sequences as well as all *Weissella rplB* sequences in CodonCode Aligner v. 9.0.1 (CodonCode Corporation, Centerville, MA, U.S.A.), and manually curated the alignments. Then, we performed Maximum Likelihood phylogenetic analysis in MEGAX (Kumar et al., 2018), utilizing the GTR model with gamma-distributed rates and invariant sites (G + I), five discrete gamma categories, and one thousand bootstrap replicates. Trees were visualized using TreeGraph (Stöver & Müller, 2010).

### Earth Microbiome Project data exploration

To more broadly understand the distribution of *Weissella,* the most abundant microbe in our dataset, we explored distributions of sequences classified as *Weissella* in the Earth Microbiome Project dataset (Thompson et al., 2017), representing the largest microbial community survey across environments to date. Toward this end, we first obtained the file containing reference-based OTUs classified using the SILVA database, emp_cr_silva_16S_123.qc_filtered.biom, from the “Data and Code” section of the project website, https://press.igsb.anl.gov/earthmicrobiome/. Using custom commands and scripts we obtained all samples containing OTUs classified as *Weissella* and as *Weissella ceti.* These samples were later classified according to study and sample source to analyze *Weissella* distribution across environments. Proportions of samples where *Weissella* was present, and proportions of *Weissella*-positive samples where at least some genotypes were classified as *W. ceti*, were compared across habitats using Chi-square test implemented in R version 3.6.3. Because of highly uneven numbers of samples from different habitats that were characterized, for those comparisons we excluded those habitats that were represented in the dataset poorly - where the “expected” number of positives, assuming even *Weissella* distribution across habitats, was below 10.

## Results

### Species identification and phylogeny

We obtained high-quality *COI* sequences for all ant workers and larvae, for 89 of the 96 myrmecophilous beetles, and for 5 out of 6 free-living beetle specimens. In most of the remaining cases, the noisier barcode sequences closely resembled some of the clean sequences, allowing for specimen classification. For all but eight myrmecophilous specimens, DNA barcodes clustered into clades that closely matched morphologically delimited bins (Table S1). One of these eight samples had a DNA barcode that was too noisy to allow classification by *COI* sequence. The seven remaining sequences matched either amphipod or nematode sequences, presumed food or parasites. These specimens were classified based on morphology (Table S1 and S3).

Based on these data, we concluded that our collection comprised 13 myrmecophile species (Fig. 1B, Fig. 1C and Supplementary appendix) from three families: Staphylinidae, Ptiliidae, and Histeridae. The four free-living beetle species belonged to the family Staphylinidae. High-quality, unambiguous sequences of representative specimens from each of the species were used for the phylogenetic reconstruction of the relationships among myrmecophiles (Fig. 1B, Supplementary appendix). Overall, by combining barcoding and morphological information we obtained the critical framework for interpreting microbiota similarity and symbiont distributions across multi-species ant myrmecophile communities. However, the identification of species was challenging, and some of them might represent undescribed species. In Supplementary Appendix, we explain how we assigned taxonomic IDs through the comparison of *COI* barcode sequences and morphological features with myrmecophiles of a more comprehensively studied *E. burchellii foreli* subspecies.

### Microbial community composition

The 16S rRNA amplicon sequencing dataset comprised 135 libraries, with a median of 20,346 reads (range 1,367-69,222). After denoising and decontamination, we obtained 5,603 microbial genotypes (zOTUs), which were grouped into 1,656 Operational Taxonomic Units (OTUs) at 97% identity. Of those, 28 OTUs, comprising a summed average of 85% of filtered reads per sequence library, were selected for more detailed analysis, according to the criteria specified in the Methods (Fig. 2).

Results for *Eciton burchellii* workers – based on a partially overlapping sample set – resembled prior observations (Łukasik et al., 2017). Their microbiota were dominated by Unclassified Firmicutes and Unclassified Entomoplasmataceae – two bacterial clades identified as specialized and stable residents of army ant gut habitats, with no close relatives identified outside of doryline ants (Funaro et al., 2011; Łukasik et al., 2017). The OTUs corresponding to these bacteria comprised, on average, 35% and 32% of *E. burchellii* worker sequence libraries, being present in 19 or 22 of the 23 characterized workers, respectively. However, both these bacteria were rare across larvae (Fig. 2). Further, reads assigned to these OTUs were scarce across myrmecophiles, generally occurring at a level expected from cross-contamination in MiSeq lanes dominated by army ant libraries (Illumina, 2017). Few other symbionts were abundant/common within or among *E. burchellii* worker libraries. Among the exceptions were *Weissella* (Lactobacillales), *Tokpelaia* (Rhizobiales), and an unclassified clade in the order Micrococcales, all found sporadically across workers.

The symbiotic microbiota of *Eciton burchellii* larvae were distinct from worker-associated microbiota, being consistently dominated by *Weissella*. Found in all ten larval specimens, the lone 97% OTU assigned to this genus accounted for 73% of reads in larval libraries, on average. The second most abundant member of the larval microbiota was *Rickettsiella* (Gammaproteobacteria: Legionellales), present in six out of ten larvae, with an average relative abundance of 10% in larval libraries (averaged across infected and non-infected specimens). The worker-dominant *Unclassified Firmicutes* was present in four individual larvae, with a low average abundance of 0.29% (range 0.029-1.42%). OTUs from *Enterococcus* and *Lactococcus* numbered among those with sporadic presence and low abundance in *E. burchellii* larvae.

While small sample sizes limit our power, we observed no clear differences between starved specimen libraries and those of their non-starved, conspecific counterparts (Fig. S2). Regardless of treatment, *Cephaloplectus mus* specimens had *Weissella* and *Lariskella* OTUs as the dominant symbionts, while *False Lomechusini sp.2* specimens had *Rickettsia* as the most abundant OTU, followed by *Weissella* (Fig. 2).

Microbiota from myrmecophilous beetles showed a higher variability. Sequences from the alphaproteobacterial genus *Rickettsia* comprised the most dominant bacterial 97% OTU, representing on average 15% of the total reads, and present at relative abundance >0.1% in 46 of 96 libraries. Other abundant and widespread bacteria included the aforementioned *Weissella* OTU, comprising 14% of the total reads among all libraries, on average, and present in 51 libraries. Reads clustering within a single *Rickettsiella* OTU (the same that was found in army ant larvae) made up 6% of reads per library, on average, and were present in 34 libraries. *Wolbachia* was represented by two OTUs, OTU5 with an average abundance of 8% and presence in 15 libraries, and OTU3 – found at 12% relative abundance per library, appearing in 26 libraries. Some of these myrmecophile-associated microbes were highly species-specific. For example, the OTU classified as *Lariskella* was abundant in about half of *Cephaloplectus mus* beetles but virtually absent in all other samples. However, other microbial OTUs abundant in the dataset were observed in multiple host species. This was particularly clear for *Weissella,* occurring in all twelve myrmecophile species, in addition to army ant worker and larvae libraries. In eight of these twelve myrmecophilous species, this OTU was present with a relative abundance of 5% or higher in at least one individual. Likewise, *Rickettsiella* was present in six army ant larvae and eleven myrmecophilous species. Many other bacteria were found in more than one beetle species, but not in army ant workers or larva. Numbering among these were a 97% OTU classified as *Rickettsia*,present in seven myrmecophilous species, and two *Wolbachia* OTUs, present in seven (OTU3) and five (OTU5) myrmecophilous species. Overall, as demonstrated in the PCoA plot, *Eciton burchellii* worker communities clearly separated from those of other insects, whereas communities of *Eciton burchellii* larvae and myrmecophilous beetles were often similar (Fig. 2, Fig. S1). The apparent abundance of prey/parasite DNA in certain beetle hosts did not lead to the detection of unique, non-beetle microbiomes, and it also did not prevent the detection of more common beetle symbionts.

Some of the microorganisms detected in myrmecophilous beetles were also found in free-living beetles. Among these, a single *Enterococcus* OTU was the most prevalent, being present in five out of six specimens in the latter category, with an average abundance of 14%. This OTU was also found in six *Eciton burchellii* larvae and 41 myrmecophilous beetles from seven species. Aside from *Enterococcus, Spiroplasma* (OTU 14) was also found in free-living. specimens (n=3) and two myrmecophilous species (n=12 specimens). However, despite such trends for these broadly distributed insect associates (Paniagua Voirol et al., 2018; Russell et al., 2012), the abundant OTUs shared among army ants and myrmecophiles were generally not present in free-living, sympatric beetles, with the exception of *Rickettsiella*, found in one free-living beetle at a relative abundance of 0.19%. Free-living beetles had high Bray-Curtis dissimilarity index when compared to myrmecophiles and ants, indicating their microbiota compositions differ (Fig. S3).

### 16S rRNA genotype-level microbial associations

Genotype-level 16S rRNA (zOTU) data provided more detailed information about the diversity and distribution of the symbiont strains within and across species (Fig. 3), and helped identify likely cases of recent transfer of symbionts among host ants and their myrmecophiles. Across the 28 selected OTUs, we identified between 1 and 13 genotypes that fulfilled the abundance criteria that were likely to exclude sequencing errors and cross-contaminants. Within a single host species, we observed up to 8 different genotypes of the same OTU, but with no clear distribution pattern across colonies (Fig. 3, Table S4). Two or more distinct genotypes from a single OTU were often found in the same individual. At the same time, the same genotypes were found in different species (Table S5) in all 24 97% OTUs shared across two or more species.

**Figure 3.**
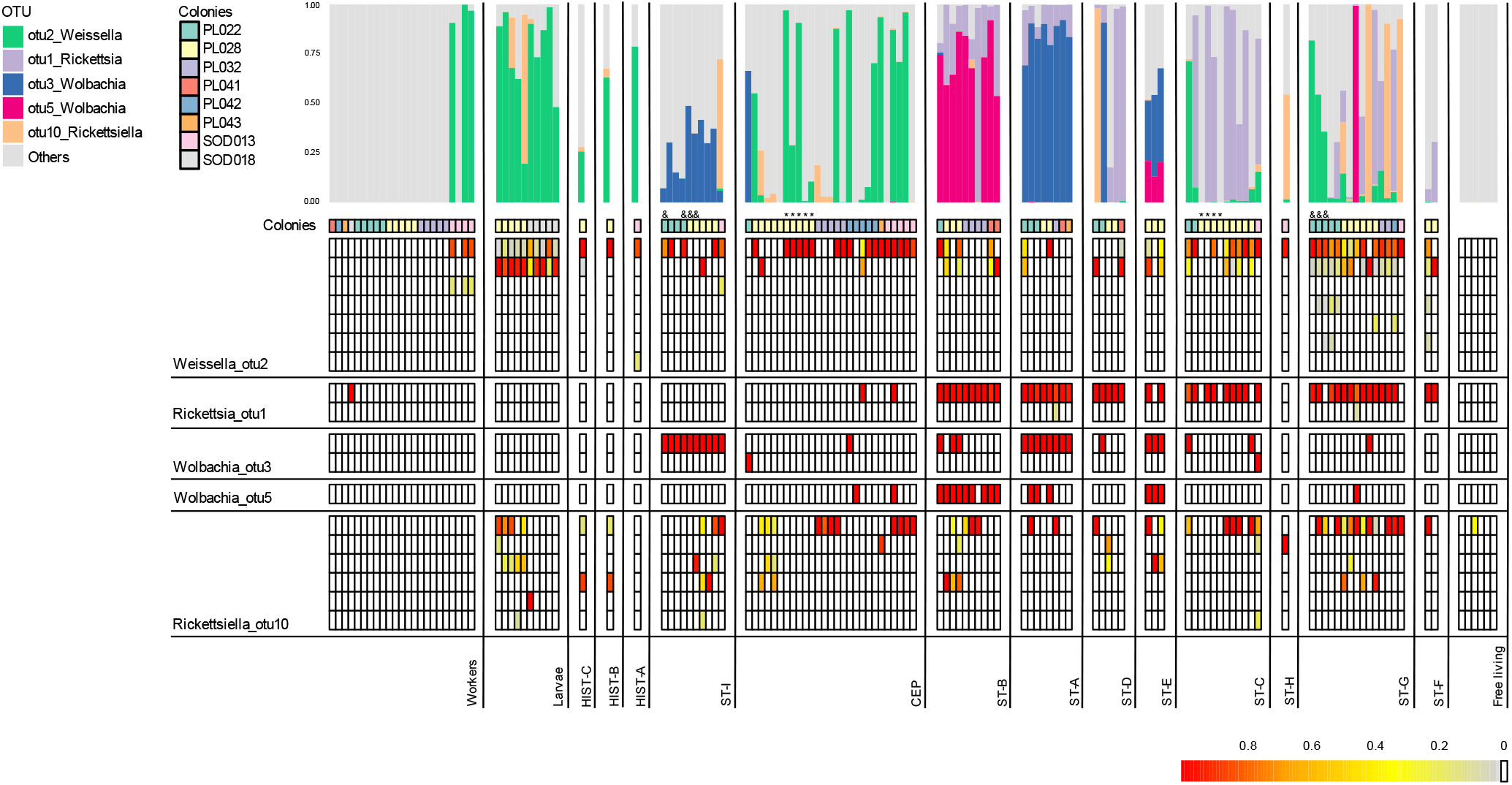
The relative abundance of 16S rRNA genotypes for selected, broadly distributed and abundant bacterial 97% OTUs. Samples are organized in columns by species, ordered based on COI ML tree, and labeled at the bottom with the abbreviation assigned in Fig.1. The bar plot shows the abundance of the OTUs relative to the total reads in each individual (columns). The heatmap presents the relative abundance of dominant genotypes within each OTU. Genotypes are shown if they have a total of at least 100 reads and a relative abundance >0.05 of the OTU in at least one sample. Asterisks mark starved specimens and ampersands mark specimens with barcode sequences matching amphipods or nematodes (see Supplementary Table S1 for details).

Genotype diversity varied among the most abundant, broadly distributed 97% OTUs (Figure 3). *Weissella* has seven detected genotypes. This was followed by the *Ricketsiella* OTU, with six genotypes, the *Rickettsia* OTU and *Wolbachia* OTU3 with two each, and by *Wolbachia* OTU5, with one. Genotype diversity also varied among host species. For example, in most myrmecophile species, we identified two *Weissella* genotypes, but *aff. Meronera* and *Ecitodonia sp*. specimens possessed up to four. In most individual insects, one genotype of *Rickettsia* was present, but in two *Ecitomorpha cf. melanotica* specimens and one aff. *Meronera* specimen, we detected additional, low-abundance genotypes. In the case of *Wolbachia* OTU3, the same genotype was detected in almost all infected insects, but one specimen of *Cephaloplectus mus* and one of *False Lomechusini sp. 2* hosted an alternative genotype. Finally, for *Rickettsiella*, most of the insects where this OTU was present harbored only one of the six genotypes, but a subset possessed two or three.

### *rplB* genotype-level diversity of Weissella

Sanger-sequencing of the protein-coding gene *rplB* for *Weissella*-positive specimens yielded 31 high-quality and unambiguous sequences. For phylogenetic analysis, we combined this dataset with sequences for 11 workers and larvae from previously characterized ant species or locations (from Łukasik et al., 2017), and sequences extracted from 25 reference genomes for other *Weissella* and other Leuconostoceae/Lactobacillales strains. The resulting maximum likelihood phylogeny was well-resolved and well-supported, providing high-resolution information on the relationships among strains. Sequences from army ants and myrmecophiles fell into two broad clades (Fig. 4). One clade included *Weissella ceti*, a species originally isolated from a beaked whale carcass (Vela et

**Figure 4.**
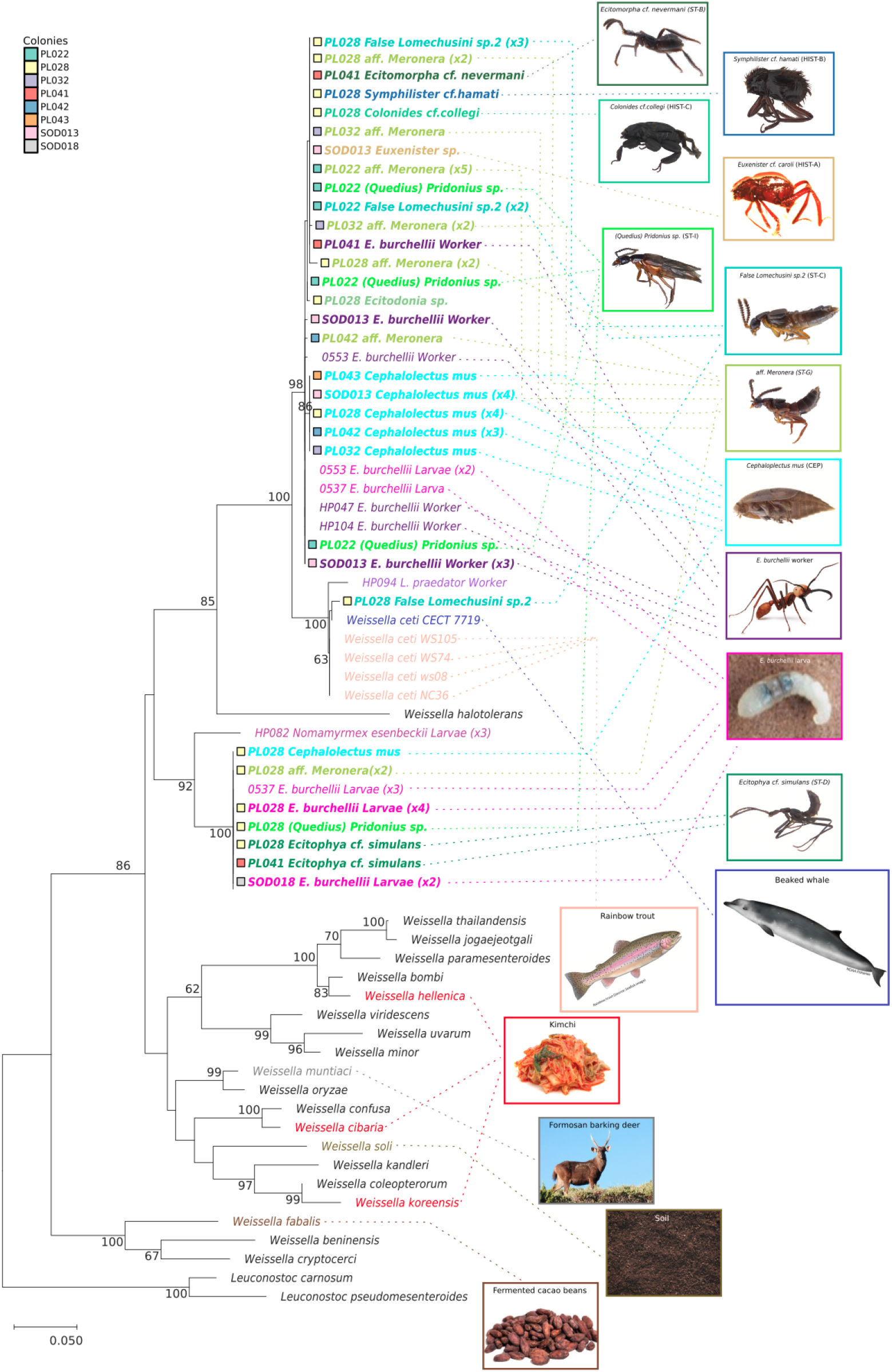
Maximum Likelihood phylogenetic tree of *Weissella* genotypes from different hosts based on the partial 510-bp sequence of the ribosomal protein L2 (*rplB*) gene. The color of the labels represents the environment or host from which the given *Weissella* isolate was derived, and it is the same color as the frame of its picture. Labels in bold letters indicate ant and myrmecophile strains from Monteverde, Costa Rica (the core samples from this project). Bootstrap values higher than 60 are shown. Identical sequences from different insects belonging to the same species and colony are represented only once in the tree, with the number of samples indicated between parenthesis next to the label.

al., 2011), and several isolates from diseased rainbow trout cultures (Figueiredo et al., 2015). While most of the newly obtained *rplB* sequences from ants and myrmecophiles were, on average, ca. 3% distinct from these previously described *Weissella* isolates, a single sequence from a myrmecophile specimen classified as *False Lomechusini sp. 2* differed by only 3 bp (0.59%) from the *W. ceti* type strain isolated from the whale. The other, divergent *Weissella* clade exclusively comprised sequences from myrmecophiles and ant larvae.

Within these two clades, the sequences from *E. burchellii* army ants and different myrmecophile species were highly similar and often identical. The *W. ceti* clade comprised sequences from nine myrmecophile species as well as army ant workers and larvae. The two most abundant *rplB* genotypes within this clade were both represented by strains from Monteverde *E. burchellii* workers and several myrmecophile species. Interestingly, one of these *rplB* genotypes was found in medium-sized *E. burchellii* workers and larvae from Venezuela. The other clade in our *rplB* phylogeny comprised a single genotype represented by sequences from four myrmecophile species and from *E. burchellii* larvae from Monteverde and Venezuela. Three identical sequences from Venezuelan *Nomamyrmex esenbeckii* army ant larvae represented a divergent genotype within that second clade.

### Earth Microbiome Project data exploration

Broad distributions of *Weissella* strains across species within army ant colonies, and their high relatedness to those from vertebrates in marine/aquatic environments, prompted us to investigate the distribution of this clade more systematically. Out of 23,828 sequence libraries from diverse habitat samples in the Earth Microbiome Project (Thompson et al., 2017), we found that 11.57% contained *Weissella*-classified OTUs, while 0.6% contained OTUs classifying as *Weissella ceti. Weissella* prevalence varied among the sampled environments (*X*^2^ (df = 14, *N* = 23730) = 4231.7, *p* < 0.001). The genus was particularly common in animal proximal gut habitats (30.52% prevalence), animal surfaces (39.82%), and surfaces from both free-living saline (52.14%) and non-saline (29.66%) categories. On the other hand, *Weissella* were entirely absent from the few sequenced “free living hypersaline” samples (Table 1, Table S7). In over half of the samples where *Weissella* was present, its relative abundance was >1%, arguing against widespread contamination or similar artifacts. Looking more specifically at the distributions of *Weissella ceti,* we found OTUs from this taxon in nearly all sampled environments. Across all environments, samples with *W. ceti* comprised <10% of those with Weissella genus, with no significant differences in *W. ceti / Weissella* ratio across environments (*X*^2^ (df = 4, *N* = 2367) = 4.0859, *p* = 0.40). In relatively few samples *W. ceti* made a substantial share (>1%) of the microbial community (Table 1, Table S7).

**Table 1.**
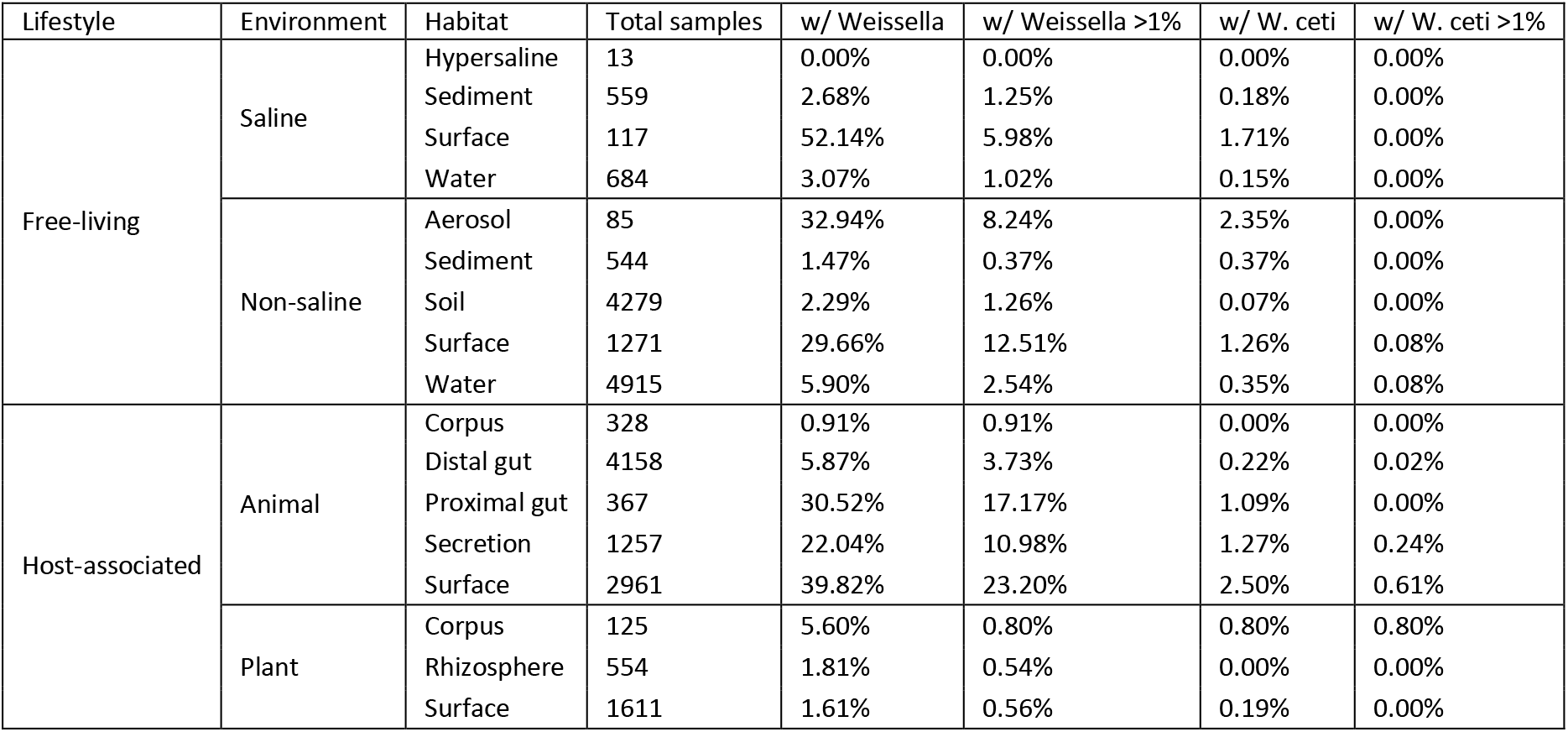
*Weissella* genus and *Weissella ceti* distribution across multiple habitats. Samples included in the Earth Microbiome Project were classified at three levels according to their source. Percentages refer to the share of samples in each “Habitat” category with any signal of *Weissella* or *W. ceti,* or where their relative abundance is at least 1%.

## Discussion

We have shown that phylogenetically distant and biologically different insects living together within army ant colonies share substantial portions of their microbiota: they frequently harbor microbes with identical genotypes at the V4 region of the 16S rRNA gene and, for *Weissella*, also at a portion of the *rplB* gene. We also found an overlap at the *Weissella* 16S-V4 and *rplB* genotype level between different *Eciton burchellii parvispinum* colonies from Costa Rica and *Eciton burchellii foreli* from Venezuela - separated by ca. two thousand kilometers and an estimated four to seven million years of evolution (Łukasik et al., 2017; Winston et al., 2017). As a reference, the specialized and putatively worker-to-worker-transmitted Firmicutes and Entomoplasmatales symbionts commonly differ among these ant colonies by about 1% within the same *rplB* gene (Łukasik et al., 2017). These patterns strongly suggest that some microbes, and *Weissella* in particular, are shared across frequently interacting species, and across geographically distant colonies of a species, at relatively short timescales. A plausible explanation of the observed patterns is its much broader environmental distribution, combined with regular transmission across host organisms or acquisition from the environment. The high similarity of ant and myrmecophile symbionts to strains isolated from phylogenetically and physiologically dissimilar animals from completely different environments, including beaked whale and rainbow trout (Figueiredo et al., 2015; Vela et al., 2011), supports the latter explanation. Combined with data on the broad distribution of *Weissella* across diverse global environments, our work highlights the versatility of some microbial clades and strains and the extent and likely significance of microbial strain transmission at multiple scales.

### Army ant workers and larvae differ substantially in their microbial community composition

The gut microbiota of Neotropical army ant workers are dominated by two clades of specialized, ancient bacterial symbionts, Unclassified Firmicutes and Unclassified Entomoplasmatales. As shown before, both were abundant in almost all workers of the most comprehensively sampled species, *E. burchellii* (Funaro et al., 2011; Łukasik et al., 2017; Mendoza-Guido et al., 2022, in press). However, as demonstrated here, these two specialized bacteria appear quite rare outside of workers. In larvae, they comprise less than 1% of the reads on average, and in myrmecophiles, they do not generally exceed levels expected from cross-contamination through index swapping during sequencing (Illumina, 2017).

In contrast, the microbial communities of army ant larvae are dominated by bacteria that appear much less specialized than those of adults. In particular, their dominant symbionts, *Weissella,* and to a lesser extent, *Rickettsiella*, are widespread, both in army ant communities and in other organisms. The consistent presence and high relative abundance of these microbes in *E. burchellii* from Costa Rica, but also the presence of closely related *Weissella* in *E. burchellii* and *Nomamyrmex* army ant larvae from Venezuela, suggests persistent association, and likely importance, in larval biology. In other social Hymenoptera, larval microbiota also differ from those of adults. For example, in *Cephalotes* turtle ants, larval microbiota can change substantially over the course of development (Hu et al., 2021), exhibiting a consistent successional pattern of unknown functional importance. Bacteria can play important roles in larvae of other social Hymenoptera. For example, a novel lactic acid bacterium was shown to inhibit the growth of the pathogen *Paenibacillus larvae* in honeybee larvae (Forsgren, et al., 2010). More systematic sampling and surveys of larval instars would clarify the significance of microbes in army ant developmental biology.

### Microbial sharing among army ant colony members

The striking degree of overlap in the bacterial associations among different myrmecophile species, and between myrmecophiles and army ant larvae, suggest extensive microbial sharing (Fig. 2). In particular, *Weissella* and *Rickettsiella*, two dominant larval associates, were both present in more than half of the sampled beetle species, in a large share of specimens. However, both were uncommon in workers; *Weissella* was present in only some specimens, and *Rickettsiella* was never detected. The identity of 16S rRNA and (in case of *Weissella*) *rplB* sequences among strains from different hosts conflicts with their strong specialization on a particular host species and instead, is highly suggestive of their ongoing or recent horizontal transmission. At the same time, closely related strains found within army ant colonies may differ in their level of host-specificity, and likely other biological characteristics. Of interest in this regard is the finding that one of the myrmecophile species, *Cephaloplectus mus*, consistently and uniquely hosted *Weissella* strains with about 0.5%*rplB* gene nucleotide sequence divergence, compared to strains that colonized a more diverse range of species (Fig. 4).

The same patterns seem to apply to most other abundant bacterial OTUs in our dataset. Several 16S rRNA genotypes of facultative endosymbionts such as *Wolbachia* and *Rickettsia* are also broadly distributed across different myrmecophile species. However, it is important to emphasize that bacterial strains identical at a 253-bp fragment of 16S rRNA may still be separated by millions of years of evolution, and differ dramatically in genome contents and the range of functions (Ochman et al., 1999). Unfortunately, we currently lack resolution to resolve the relationships among strains detected in different myrmecophile species, and cannot reach precise conclusions on the recency of the interspecific symbiont transmission. While it is likely to be ongoing in many cases, whole-genome comparison among isolates from different host species would provide the ultimate evidence.

### Army ant colonies as arenas for interspecific exchange of microbial symbionts

It appears that the extensive and close interactions among army ants and their diverse myrmecophilous beetles facilitate microbial strain transmission across species, resulting in the broad host distribution of the dominant microbial clades. While OTUs found in myrmecophiles and ants were also present in free-living beetles, the OTUs that we identified as being shared among the community, such as *Weissella* and *Rickettsia* were not (Fig. 2, Fig. 3). This is also evidenced when comparing the microbial composition similarity among samples grouped by species, where we see that Bray-Curtis dissimilarity indexes for free-living beetles are generally high, indicating low similarity among the libraries (Fig. S3). Thus, with their hundreds of associated insect species, army ant colonies appear to serve as excellent arenas for interspecific microbial symbiont exchange. The transmission is likely shaped by diverse interactions among ants and myrmecophile species, with some being bivouac inquilines while others are only present in raids or refuse deposits. *Ecitophya* and *Ecitomorpha* species, for example, can practice grooming (von Beeren et al., 2018). Similarly, in velvety tree ant (*Liometopum occidentale*) communities, ants and myrmecophiles show similarities in their microbiota composition, depending on the level of interaction they show (Perry et al., 2021). As myrmecophiles can be found in the nests of practically every ant species (Danoff-Burg, 2008; Kronauer & Pierce, 2011), the study of these relationships and their influence on the microbiota of the species involved could be of great importance to understand microbe sharing among insects.

The mechanisms of transmission among ant workers, larvae, and their associated beetles are likely to vary substantially among bacterial symbiont categories (Perreau & Moran, 2021). Bacteria that form “open” symbioses, which include most gut symbionts, are commonly acquired from or through the environment - creating opportunities for different cohabiting insects to acquire the same microbes from the same sources within a colony. Sharing food sources, predation, grooming, and other social interactions create opportunities for direct transfer of microbes from one host insect to another. In the cases of *Weissella* and *Rickettsiella*, it is tempting to assume that larvae are the primary sources of these microbes for other insects within a colony. On the other hand, in *Eciton burchellii* army ants, batches of larvae are synchronized and separated by periods when no larvae are present within colonies, preventing direct transmission from older to younger larvae (Kronauer, 2020). Given the scarce presence of these two microbes in workers, it becomes, alternatively, tempting to postulate beetles as the source of *Weissella* for new generations of larvae, despite limited evidence for direct interactions between larvae and most myrmecophile species. However, studies including other castes within the colonies are necessary to confirm this. Whichever the ultimate source, through their close integration into colony biology, many myrmecophiles are plausibly exposed to similar microbial inocula as those encountered by army ant larvae, and it seems likely that inoculation is not unidirectional.

Interspecific transmission of facultative endosymbionts such as *Wolbachia* and *Rickettsia* is likely more complicated than that of gut symbionts, but still probably facilitated within large colonies inhabited by multiple potentially suitable hosts. To establish a novel infection, heritable endosymbionts need to be physically transferred from body fluids of one insect to another, avoid the immune system, establish means of transmission to host reproductive tissue and across generations, and affect host fitness in ways that would prevent the rapid clearing of the infection by natural selection (Bright & Bulgheresi, 2010). It can be argued that at least the first step in the process - opportunity for acquiring the new infection - is facilitated among species that live closely together and interact frequently, and are exposed to shared pools of external parasites, or perhaps preying on each other (Ahmed et al., 2015; Clec’h et al., 2013). *Wolbachia* has been found in extracellular environments of attine ants (Tolley et al., 2019), *Drosophila melanogaster* (Pietri et al., 2016), and *Nasutitermes arborum* termites (Diouf et al., 2018), among others, and while the viability of the cells was not established, this might facilitate their transmission. This could also be true for other microbes thought to exist strictly as intracellular symbionts. There is also evidence that the environment insects live in might act as a reservoir of some microbes, with the *Wolbachia* signal being detected in plant matter and fungi (Li et al., 2017). However, other studies show that cohabitation does not always result in microbiota similarities, as is the case of some ant species and their trophobiont mealybugs and aphids (Ivens et al., 2018).

### Broad distributions of bacterial clades that infect ants and myrmecophiles

Several of the microbes abundant in army ant colonies are distributed much more broadly. *Wolbachia* is estimated to infect approximately half of all insect species, many nematodes, and other invertebrates (Kaur et al., 2021). Likewise, *Rickettsia* infects diverse insects, often forming persistent associations - although some strains are plant and vertebrate disease agents vectored by arthropods (McGinn & Lamason, 2021). *Rickettsiella* is also known from diverse arthropods, although the nature of these associations is often unclear (Zchori-Fein & Bourtzis, 2012).

In contrast, the genus *Weissella* is not well-known as an insect associate. Some of the named species were originally isolated from insects (*Weissella bombi* from a bumblebee, *Weissella cryptocerci* from a cockroach - Heo et al., 2019; Praet et al., 2015), others from vertebrates (e.g., *Weissella confusa*, human pathogen - Fairfax et al., 2014; Kamboj et al., 2015), and several live in fermented foods *(Weissella koreensis* in kimchi, *Weissella fabaris* in fermented cacao beans - Lee et al., 2002; Snauwaert et al., 2013). *Weissella’s* metabolic versatility combined with abundant opportunities for bacterial transmission across cohabiting species could explain its broad distribution in army ant colonies.

Explaining the close similarity of ant and myrmecophile microbes to previously described strains of *Weissella ceti is more* challenging. The bacterial species was originally isolated from the internal organs of a dead beaked whale, and was further reported as a pathogen of commercially raised rainbow trout in China (Vela et al., 2011), in Brazil, and North America (Castrejón-Nájera et al., 2018; Figueiredo et al., 2015). These two vertebrate species are physiologically dissimilar and inhabit completely different environments than army ants, and opportunities for direct microbial exchange among them must be extremely limited. Still, the close similarity within the *rplB* protein-coding gene between strains infecting fish, whales, ant larvae, and some myrmecophiles does indicate that the interspecific transmission, likely through a long chain of other hosts or habitats, must have occurred relatively recently. One can envision that *Weissella* routinely transmits across a wide range of hosts and environments. Both our analysis of the Earth Microbiome Project data (Thompson et al., 2017) demonstrating that the genus *Weissella*, as well as *Weissella ceti*, occur in nearly all sampled environments (Table 1) and literature data (Abriouel et al., 2015) strongly argue for the versatility of this microbial clade.

Such versatility may not be an unusual characteristic for a bacterial genus. Species from the genus *Lactobacillus*, for example, can be found in a wide range of vertebrate and invertebrate hosts and also in fermented plant and milk products (Zheng et al., 2020). Members of the *Enterobacter* and *Pseudomonas* genera can also be found in a broad spectrum of habitats, such as plants, soil, aerosol and water, in addition to being opportunistic pathogens or more commensal members of the gut microbiota of vertebrates and invertebrates (Grimont & Grimont, 2006; Silby et al., 2011). The Earth Microbiome Project has reported many other bacterial clades broadly distributed and abundant across environments, including *Bacillus, Enterobacteriaceae, and Streptococcus* (Thompson et al., 2017; https://earthmicrobiome.org/). But we are far from understanding their ecological and evolutionary relevance in different environments.

However, it seems less common for a bacterial species to be present in very different environments or hosts, especially if there is no interaction between them. With the sampling bias towards medically and agriculturally important microbes, most known multi-host bacterial species are vertebrate pathogens, such as *Vibrio cholerae* or *Yersinia pestis*, with different hosts often harboring distinct strains (Bacigalupe et al., 2019). Both pathogens are also occasionally found in the environment outside of host organisms. Another such microbe is *Pseudomonas syringae*, which can be found in plants and aerosols, as well as a pathogen of some aphid species (Alsved et al., 2018; Smee et al., 2021). We expect that broader and focused sampling of species, environments, and other samples that are not of immediate medical, agricultural, or economic importance, will reveal a much wider range of broadly distributed microbial species.

### Biological properties and fitness effects as a critical aspect of microbial transmission and distribution

Depending on the nature of the association with their hosts, symbiotic microorganisms vary in their fitness effects, transmission propensity, and evolutionary potential. Facultative endosymbionts such as *Rickettsia*, *Wolbachia, and Spiroplasma* have been traditionally regarded as reproductive parasites, but more recent research has revealed a range of functions that can clearly benefit hosts, including the biosynthesis of nutrients and protection against natural enemies (Kaur et al., 2021; Łukasik et al., 2013; Nikoh et al., 2014). Through the combination of reproductive manipulation and fitness benefits, new infections with these microbes, likely initially acquired from other species, have swept through host populations (Cockburn et al., 2013; Himler et al., 2011; Kriesner et al., 2013), with effects likely reverberating in multi-species communities (Ferrari & Vavre, 2011). At short timescales, such infections could enable rapid response and adaptation to environmental challenges, particularly relevant in the rapidly changing world of the Anthropocene(Lemoine et al., 2020). At longer timescales, they may facilitate and speed up speciation (Janson et al., 2008; Moran, 2007). The patterns and processes relevant to the distribution and transmission of these microbes are increasingly recognized as an essential component of their hosts’ biology.

For facultative symbionts, represented by *Weissella* and *Rickettsiella* species, we are only starting to unravel their distributions, the spectra of their functional diversity, details of their associations with host organisms, and ecological and evolutionary significance. With highly fragmented and biased data, we are far from understanding any of these processes in non-model organisms, including millions of insect species that have not yet been formally described (Adis, 1990; Stork, 2018), and which are increasingly threatened by extinction as climate change and other anthropogenic disturbances intensify (Raven & Wagner, 2021). Army ant colonies serve as convenient arenas for interspecific exchange of various microbes, and it is likely that such microbial exchange is common in other environments, for example, where animals share food resources (Stahlhut et al., 2010), or in predator-prey interactions (Clec’h et al., 2013; Kennedy et al., 2020). To fully understand the processes and patterns related to microbial transmission across species, their dynamics, and significance, it is clear that future broad surveys of microbiota across diverse wild insect communities will need to include a comprehensive analysis of their ecology and interactions with other organisms.

## Acknowledgments

We thank Minoru Moriyama, Alfred F. Newton, and Mariana Chani-Posse for help with specimen identification and Szymon Drobniak for statistical advice. We also thank the Symbiosis Evolution Group at Jagiellonian University, especially Diego Castillo-Franco, Sandra Åhlén Mullio and Junchen Deng, for their participation in discussions about the results and how to present them. The project was supported by the U.S. National Science Foundation grant 1050360 to J.A.R., Polish National Agency for Academic Exchange grant PPN/PPO/2018/1/00015, and Polish National Science Centre grant 2018/31/B/NZ8/01158 to P.Ł.

## Data Accessibility

Sanger sequences were deposited in GenBank (Accession nos OP850275-OP850288). Amplicon sequencing data were deposited in NCBI Sequence Read Archive (BioProject: PRJNA900236; 16S rRNA data: SAMN31683713-SAMN31683857). Colony and insect details are provided in the Supplementary Tables S1 and S2. In Tables S1 and S6, we listed the above accession numbers by individual. All samples were collected in accordance with national and international laws; collection permit numbers include 122-2009, 192-2012 and R-009-2014-OT-CONAGEBIO (Costa Rica).

## Author Contributions

P.Ł., J.A.R. and C.V. designed the research. P.Ł., D.J.K., S.O.D. and C.v.B. provided specimens and identifications. P.Ł. and J.A.N. generated molecular data. C.V. and P.Ł. analysed the molecular data, C.V. and C.v.B. illustrated the results. C.V., P.Ł. and J.A.R. wrote the manuscript, and all authors contributed to revisions.

