## Supplementary Figures for "Microbial symbionts are shared between ants and their associated beetles"

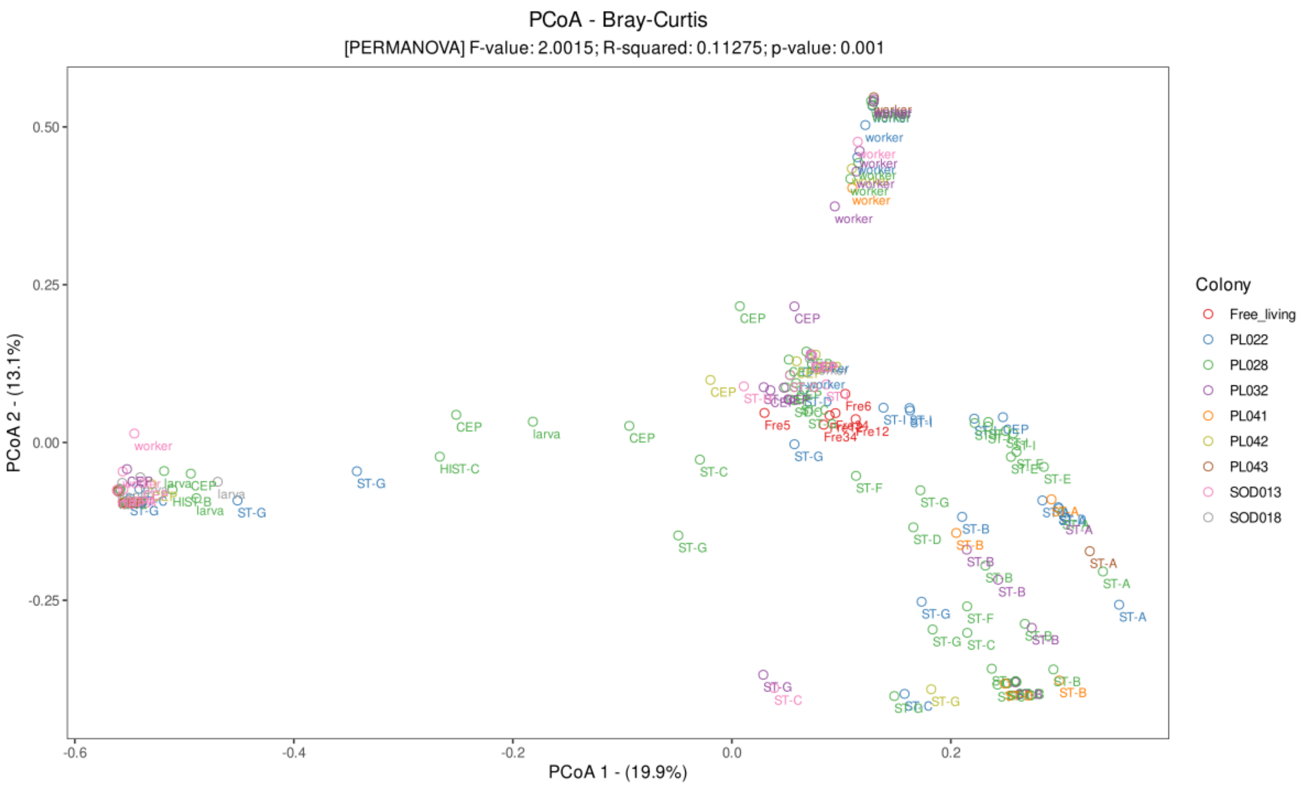


**Figure S1.** Principal coordinates analyses (PCoA) plot for *Eciton burchellii* and associated myrmecophilous beetles, based on microbial relative abundance, with 999 permutations. Labels correspond to the species name abbreviations (see Table S3), and colors represent different army ant colonies. Metrics were calculated using the Phyloseq 1.30.0 and Vegan 2.6-2 packages and visualized using ggplot2 3.4.0 using R version 3.6.3.

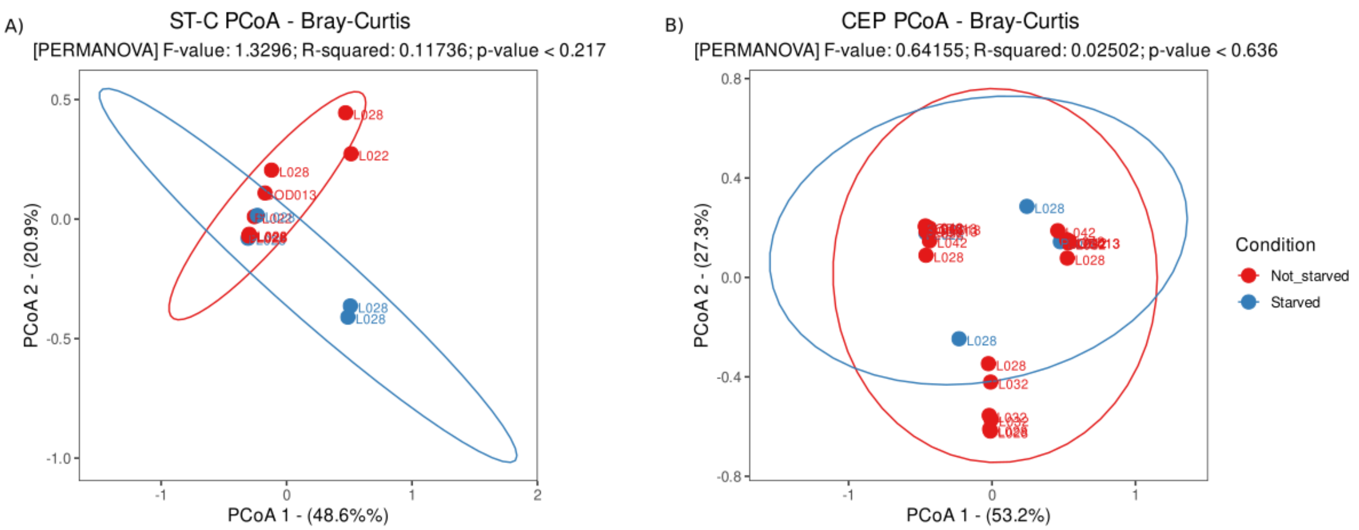


**Figure S2.** Principal coordinates analyses (PCoA) plots for specimens of two myrmecophile species, collected from two ant colonies, in which some specimens were starved prior to preservation, based on microbial relative abundance, with 999 permutations. A) PCoA of starved and non-starved False *Lomechusini sp.2* specimens. B) PCoA of starved and non-starved *Cephaloplectus mus* specimens. Labels correspond to the individual's colony. Metrics were calculated using the Phyloseq 1.30.0 and Vegan 2.6-2 packages and visualized using ggplot2 3.4.0 using R version 3.6.3.

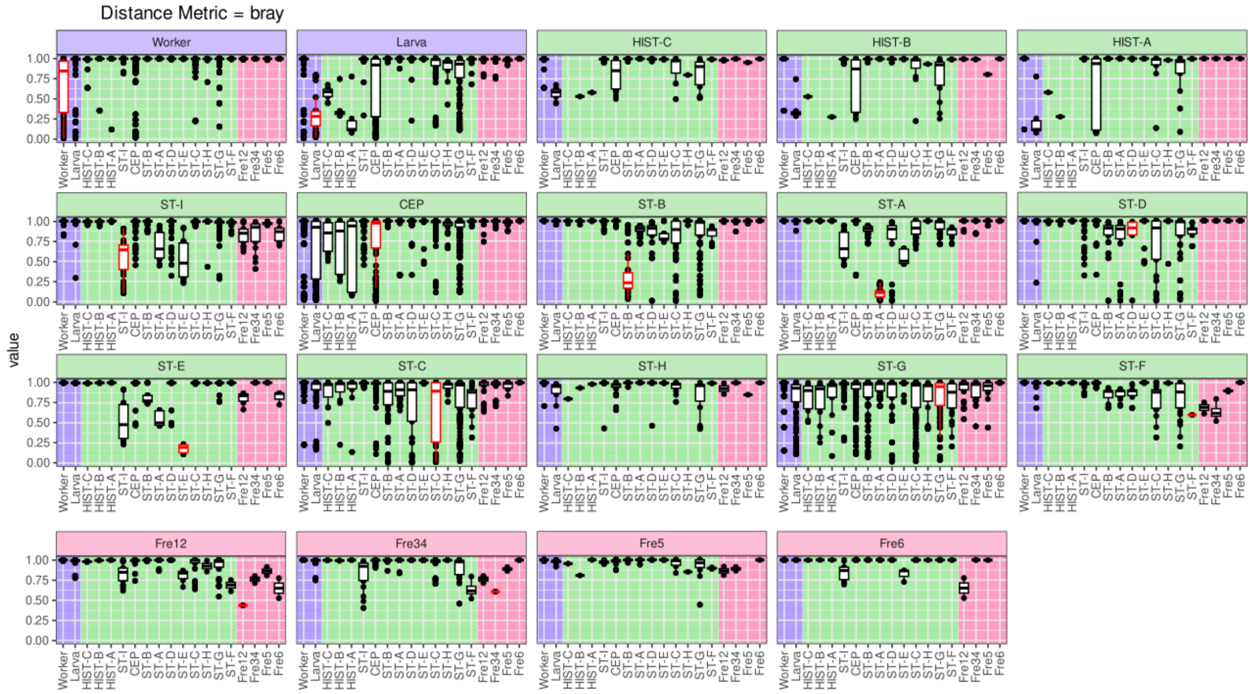


**Figure S3.** Bray-curtis dissimilarity pairwise comparison of samples from each species versus the rest. Tittle and background color indicate which group the species belongs to: ants (purple), myrmecophiles (green) and or free-living insects (pink). Red box indicates results for samples belonging to the same species they are being compared. Self-comparisons were removed (when there is no red box, only one individual was analyzed for that species). Metrics were calculated using the Phyloseq 1.30.0 and Vegan 2.6-2 packages and visualized using ggplot2 3.4.0 using R version 3.6.3.
