## Supplementary appendix for "Microbial symbionts are shared between ants and their associated beetles"

### Supplemental material – Myrmecophile identification protocol

We used the images and the *COI* barcodes for identifications. The specimens themselves were not available because they were destroyed during the genetic procedures. Providing morphological and *COI* characters, the recently published inventory of *Eciton* guests of von Beeren et al. (2021) served as a baseline for identifications. We mainly relied on the *COI* barcode to distinguish between morphologically very similar species and used a combination of barcodes and the images to identify the myrmecophiles to the lowest taxonomic level possible. References to species keys are provided in von Beeren et al. (2021). Voucher images of reference specimens have been uploaded to the Barcode of Life database system (http://www.boldsystems.org/). Images of the herein studied myrmecophiles are attached below.

First of all, one main impediment of species identifications needs to be addressed. Regarding army ants, Costa Rica is divided into two main biogeographic regions, the part west of the volcanic front, the 'Chrotega block', and the part east of it (Winston, Kronauer, & Moreau, 2016). The western part is mainly inhabited by the army ant subspecies *Eciton burchellii parvispinum* Forel, 1899*,* while the eastern part is mainly inhabited by the subspecies *Eciton burchellii foreli* Mayr, 1886 (Winston et al. 2016; see also antmaps.org). It seems likely to us that these two differentially colored subspecies represent two distinct species as indicated by genetic and biogeographic data (Pérez-Espona, Goodall-Copestake, Berghoff, Edwards, & Franks, 2017; Winston et al., 2016).

In the present work, the myrmecophiles were collected from the subspecies *E. burchellii parvispinum* at Monteverde, while the recently published *Eciton* guest inventory focused on myrmecophiles from the host ant *E. burchellii foreli* collected at La Selva Biological Station (von Beeren et al., 2021). Because many army ant associated myrmecophiles are specifically associated with a single host species only (von Beeren et al., 2021), some uncertainty remains of whether certain myrmecophiles studied here represent the same, or closely related (sister) species to those myrmecophiles studied at La Selva. In highly host-specific army ant myrmecophiles such as *Euxenister caroli* Reichensperger, 1923 and the myrmecoid beetles of the genera *Ecitomorpha* and *Ecitophya* (Ivens, von Beeren, Blüthgen, & Kronauer, 2016; von Beeren et al., 2021, 2018), we consider it likely that they rather represent sister species. Some evidence favors this scenario. A recent study indicated that the two mentioned army ant subspecies harbored different, but closely related *Ecitophya* Wasmann, 1900 and *Ecitomorpha* Wasmann, 1889 species (Pérez-Espona et al., 2017). Similarly, in the phylogenetic tree below, we found a relatively high sequence divergence between the Monteverde and La Selva population in morphologically similar specimens that keyed out as the same species.

In the following we give a few examples. The specimens morphologically identified as *E. caroli* merely showed a 88% sequence similarity to those *E. caroli* specimens studied by von Beeren et al. (2021). This is a low degree of similarity for *COI* barcodes within a species, even across geographic areas. Morphologically, specimens from both populations clearly keyed out as *E. caroli*. Noteworthy, a male of this species collected at Las Cruces, CR, from an *Eciton burchellii parvispinum* colony appears to be morphologically similar but probably distinct enough to consider it a different species to those specimens collected in *E. burchellii foreli* colonies (personal communication, A. Tishechkin). Future integrative work will bring clarity regarding their taxonomic status, but this was beyond the scope of the present study. We decided to denote the specimens studied here as *Euxenister* cf. *caroli* (cf= confer; probably belongs to the identified or a close sister species).

We also encountered difficulties in assigning species names in the myrmecoid staphylinid beetles of the genus *Ecitomorpha*. Kistner & Jacobson synonymized three *Ecitomorpha* species so that the genus formally contains a single species - *E. arachnoides* Wasmann 1889 (Kistner & Jacobson, 1990). However, it was previously demonstrated that this taxon represents a species complex (Pérez-Espona et al., 2017; von Beeren et al., 2018). Based on *COI* barcode clustering, we discovered two distinct *Ecitomorpha* species. One of them matched morphologically closest to the species *Ecitmorpha nevermanni* Reichensperger 1935 (see also von Beeren et al. 2018 for species characters) and the *COI* barcodes additionally matched closest to this species (von Beeren et al. 2021; 96% sequence similarity). We thus denoted this species as *Ecitomorpha* cf. *nevermanni*. With only 90% of base pairs similarity to the species *Ecitomopha* cf. *breviceps* - the closest match in the reference DNA barcode database - we remain uncertain in identifying the second detected *Ecitomorpha* species based on DNA barcode data. The species showed morphological characters that best match to the species description of *Ecitomorpha melanotica* Mann, 1926 (see Reichensperger, 1933). We did not study *E. melanotica* type material, leaving uncertainties in species identification in this challenging group. However, similar to the collection of the present work this species was collected with *Eciton burchellii parvispinum* at Monteverde (Akre & Rettenmeyer, 1966). As noted by several authors (Akre & Rettenmeyer, 1966; Kistner & Jacobson, 1990), *Ecitomorpha* specimens match the color of their *E. burchellii* subspecies in that *E. melanotica* is darker than the *Ecitomorpha* specimens collected in Central America with *E. burchellii foreli*. The specimens analyzed here were indeed darker, and we thus decided to denote the herein studied species as *Ecitomorpha* cf. *melanotica*. As stated previously (von Beeren et al., 2018), this interesting group of ant ant-mimicking beetles is in urgent need of a taxonomic revision covering the entire biogeographic range of these guests. Unfortunately, this is true for many myrmecophile groups of Neotropical army ants, which puts limits on accuracy of species identifications. For the present work this means we cannot unambiguously identify species in many cases and express this with the terms 'cf.' (confer) and 'aff.' (affinis; having similarities but is not identical). However, by providing voucher DNA barcodes as well as voucher images of the different myrmecophiles, our study will hopefully allow future work to clarify the uncertainty in the present work’s species identifications.

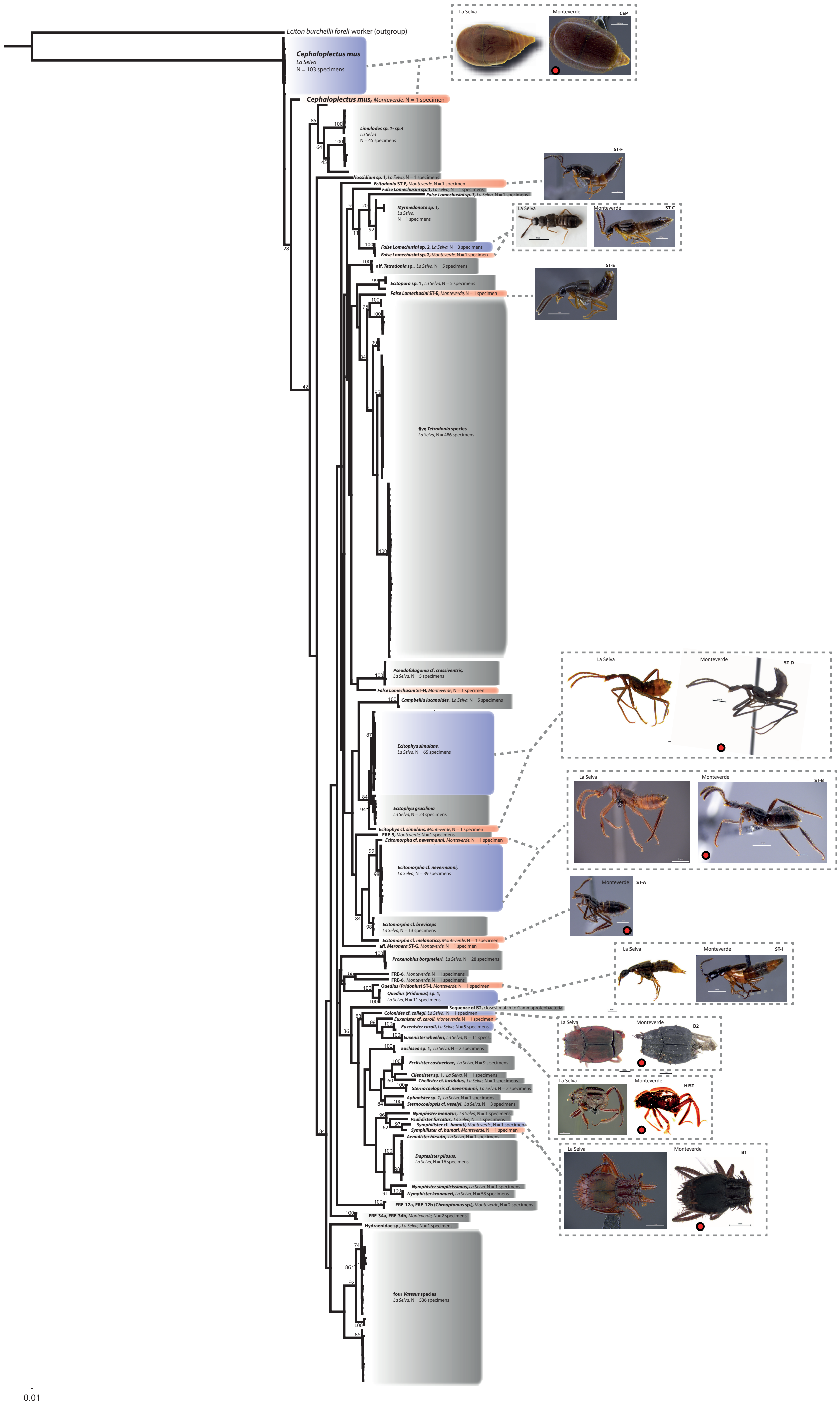

Eciton burchellii foreli worker (outgroup)
Cephaloplectus mus
La Selva
N = 103 specimens
Cephaloplectus mus, Monteverde, N = 1 specimen
Limulodes sp. 1- sp.4
La Selva
N = 45 specimens
Nossidium sp. 1, La Selva, N = 1 specimens
Ecitodonia ST-F, Monteverde, N = 1 specimen
False Lomechusini sp. 1, La Selva, N = 1 specimens
False Lomechusini sp. 3, La Selva, N = 1 specimens
Myrmedonota sp. 1,
La Selva,
N = 1 specimens
False Lomechusini sp. 2, La Selva, N = 3 specimens
False Lomechusini sp. 2, Monteverde, N = 1 specimen
aff. Tetradonia sp. , La Selva, N = 5 specimens
Ecitopora sp. 1 , La Selva, N = 5 specimens
False Lomechusini ST-E, Monteverde, N = 1 specimen
five Tetradonia species
La Selva, N = 486 specimens
Pseudofalagonia cf. crassiventris,
La Selva, N = 5 specimens
False Lomechusini ST-H, Monteverde, N = 1 specimen
Campbellia lucanoides , La Selva, N = 5 specimens
Ecitophya simulans,
La Selva, N = 65 specimens
Ecitophya gracilima
La Selva, N = 23 specimens
Ecitophya cf. simulans, Monteverde, N = 1 specimen
FRE-5, Monteverde, N = 1 specimens
Ecitomorpha cf. nevermanni, Monteverde, N = 1 specimen
Ecitomorpha cf. nevermanni,
La Selva, N = 39 specimens
Ecitomorpha cf. breviceps
La Selva, N = 13 specimens
Ecitomorpha cf. melanotica, Monteverde, N = 1 specimen
aff. Meronera ST-G, Monteverde, N = 1 specimen
Proxenobius borgmeieri, La Selva, N = 28 specimens
FRE-6, Monteverde, N = 1 specimens
FRE-6, Monteverde, N = 1 specimens
Quedius (Pridonius) ST-I, Monteverde, N = 1 specimen
Quedius (Pridonius) sp. 1,
La Selva, N = 11 specimens
Sequence of B2, closest match to Gammaproteobacteria
Colonides cf. collegi, La Selva, N = 1 specimen
Euxenister cf. caroli, Monteverde, N = 1 specimen
Euxenister caroli, La Selva, N = 5 specimens
Euxenister wheeleri, La Selva, N = 11 specs.
Euclasea sp. 1, La Selva, N = 2 specimens
Ecclisister costaericae, La Selva, N = 9 specimens
Clientister sp. 1, La Selva, N = 1 specimens
Cheilister cf. lucidulus, La Selva, N = 1 specimens
Sternocoelopsis cf. nevermanni, La Selva, N = 2 specimens
Aphanister sp. 1, La Selva, N = 1 specimens
Sternocoelopsis cf. veselyi, La Selva, N = 3 specimens
Nymphister monotus, La Selva, N = 1 specimens
Psalidister furcatus, La Selva, N = 1 specimens
Symphilister cf. hamati, Monteverde, N = 1 specimen
Symphilister cf. hamati, Monteverde, N = 1 specimen
Aemulister hirsuta, La Selva, N = 1 specimens
Daptesister pilosus,
La Selva, N = 16 specimens
Nymphister simplicissimus, La Selva, N = 1 specimens
Nymphister kronaueri, La Selva, N = 58 specimens
FRE-12a, FRE-12b (Chroaptomus sp.), Monteverde, N = 2 specimens
FRE-34a, FRE-34b, Monteverde, N = 2 specimens
Hydraenidae sp., La Selva, N = 1 specimens
four Vatesus species
La Selva, N = 536 specimens
0.01
La Selva
Monteverde
CEP
ST-F
ST-C
ST-E
ST-D
ST-B
ST-A
ST-I
B2
HIST
B1

| Species LIM |
| --- |
| 22-LIM-2 |
| 28-LIM-1 |
| 28-LIM-2 |
| 28-LIM-3 |
| 28-LIM-S1 |
| 28-LIM-S2 |
| 28-LIM-S3 |
| 32-LIM-1 |
| 32-LIM-2 |
| 32-LIM-3 |
| 32-LIM-4 |
| 42-LIM-3 |
| 42-LIM-5 |
| 43-LIM-1 |
| SOD013-LIM-2 |
| SOD013-LIM-3 |
| SOD013-LIM-4 |
| SOD013-LIM-6 |
| SOD013-LIM-7 |

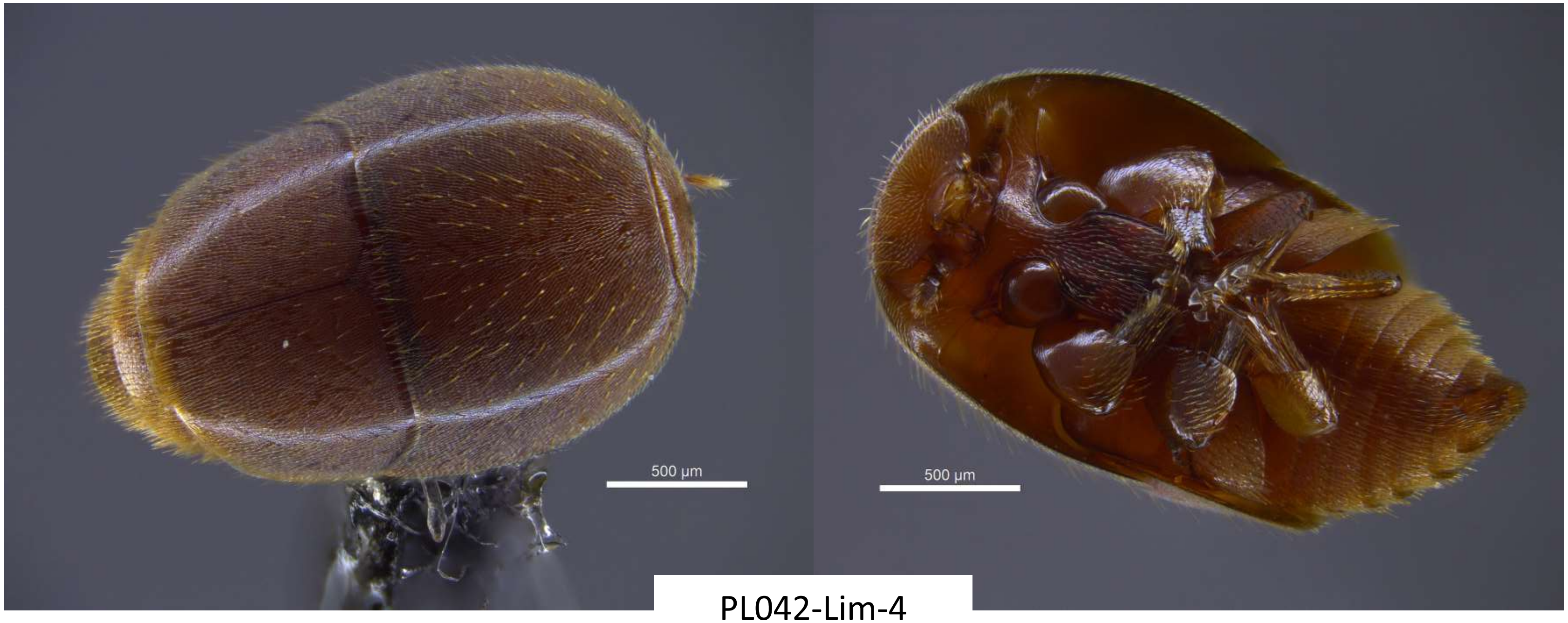


PL042-Lim-4

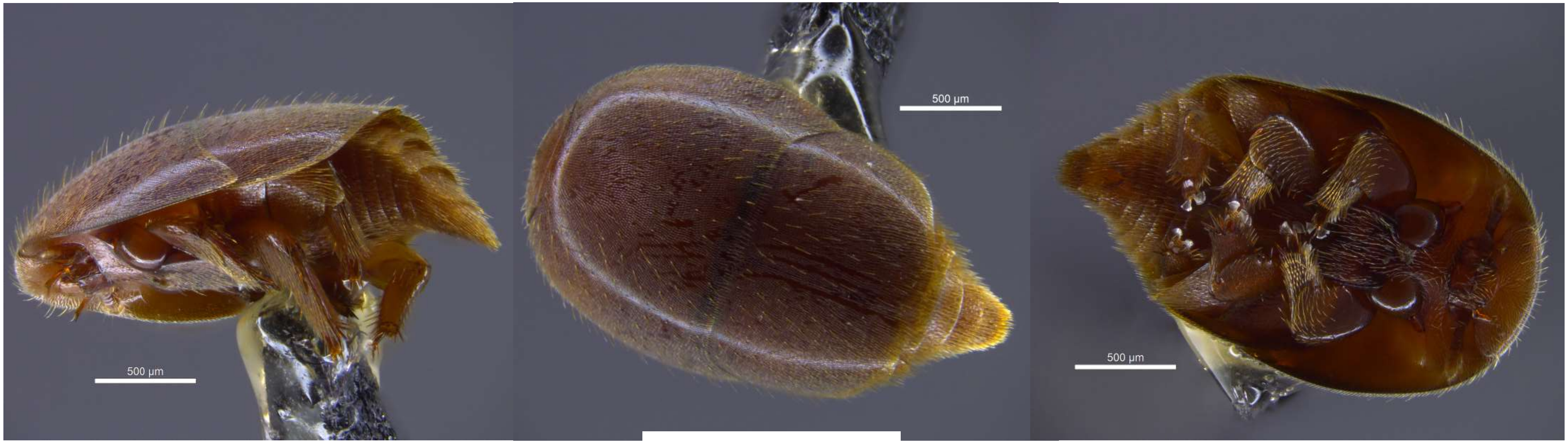


PL028-Lim-4

| Species ST-A |
| --- |
| PL022-ST6-1 |
| PL022-ST6-2 |
| PL022-ST6-3 |
| PL028-ST6-2 |
| PL028-ST6-5 |
| PL032-ST6-4 |
| PL041-ST6-4 |
| PL043-ST6-1 |

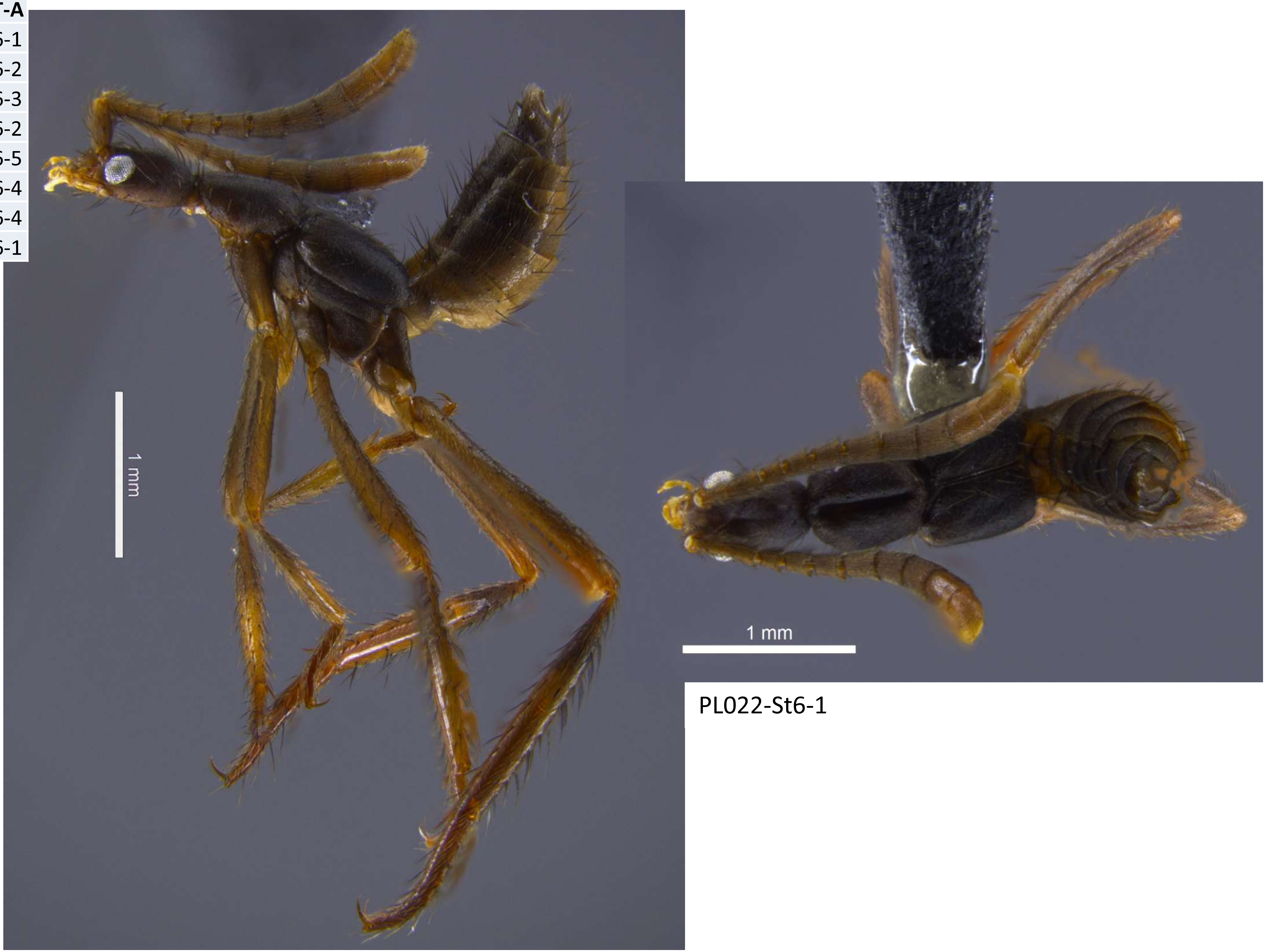


PL022-St6-1

| Species ST-A |
| --- |
| PL022-ST6-1 |
| PL022-ST6-2 |
| PL022-ST6-3 |
| PL028-ST6-2 |
| PL028-ST6-5 |
| PL032-ST6-4 |
| PL041-ST6-4 |
| PL043-ST6-1 |

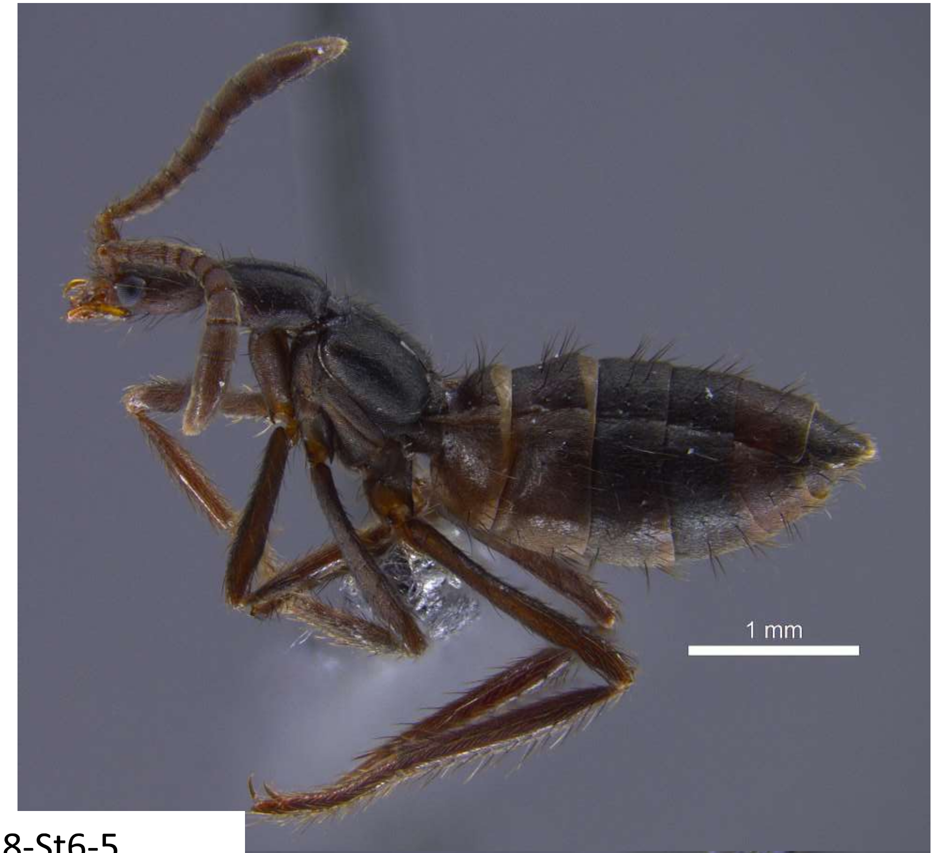


PL028-St6-5

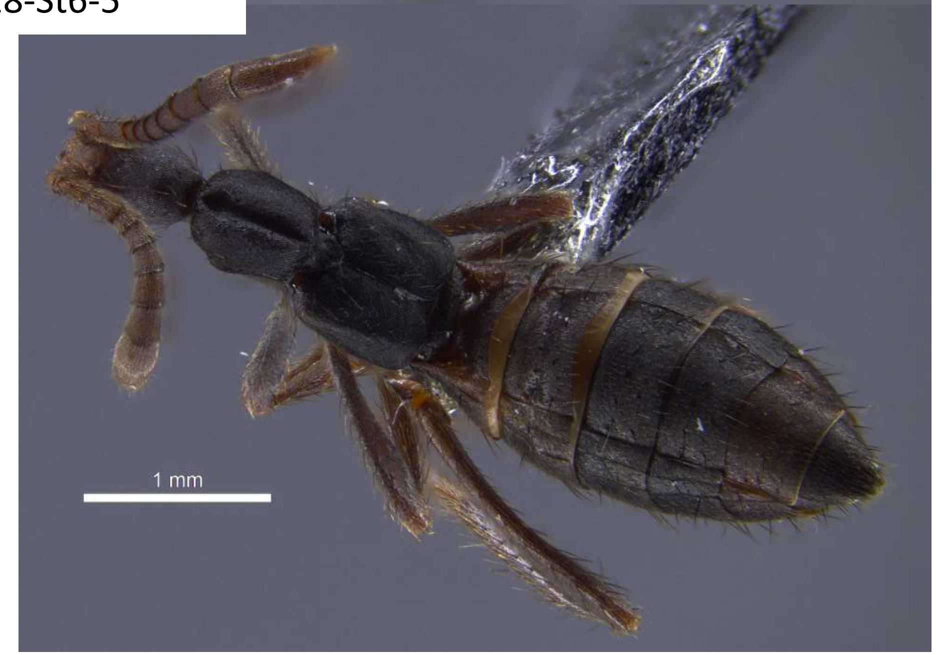


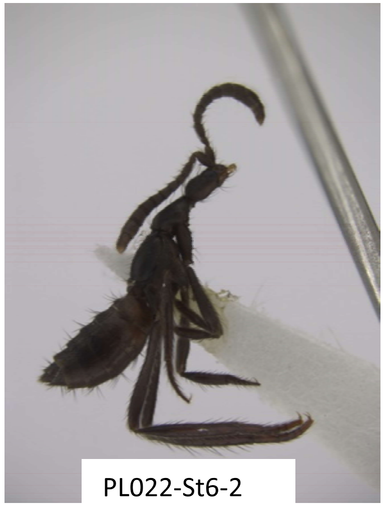


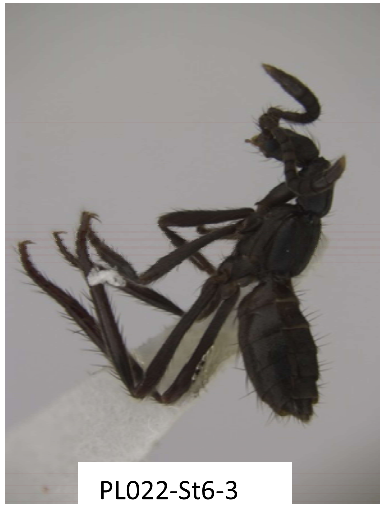


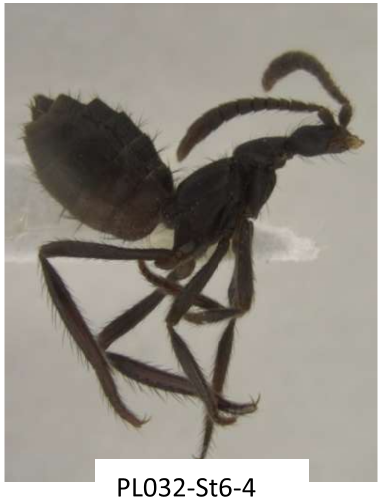


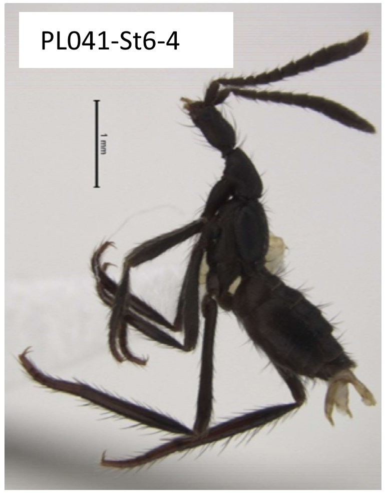


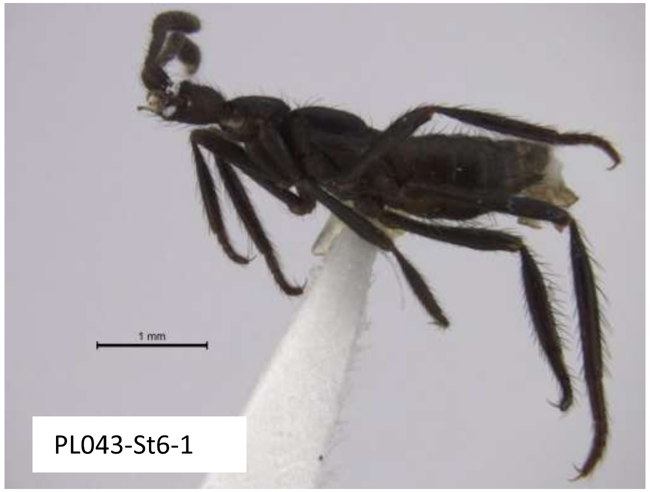

| Species ST-B |
| --- |
| PL022-ST6-4 |
| PL028-ST6-1 |
| PL028-ST6-3 |
| PL028-ST6-4 |
| PL032-ST6-1 |
| PL032-ST6-2 |
| PL032-ST6-3 |
| PL032-ST6-5 |
| PL041-ST6-2 |
| PL041-ST6-3 |

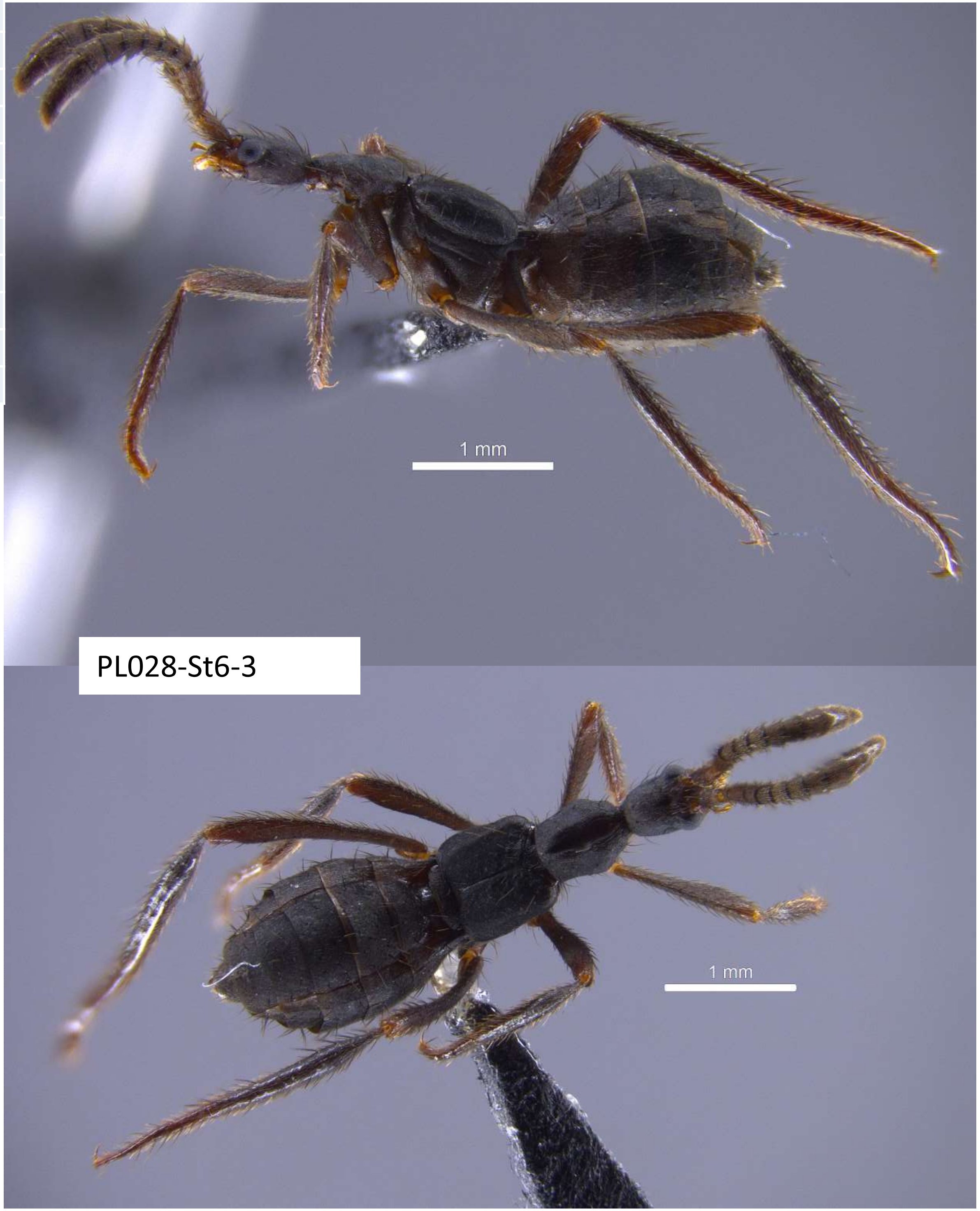


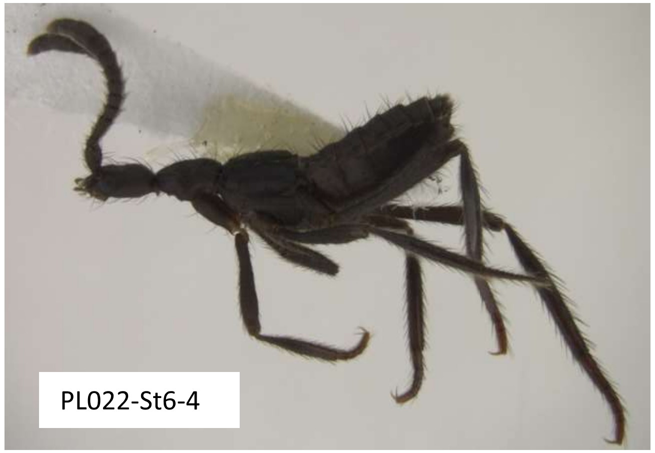


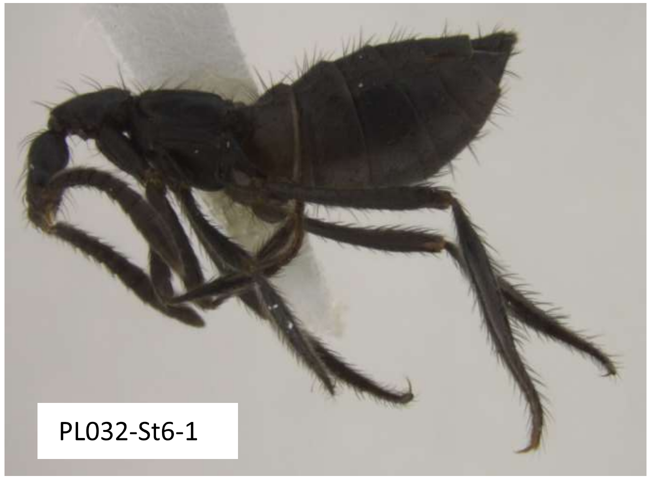


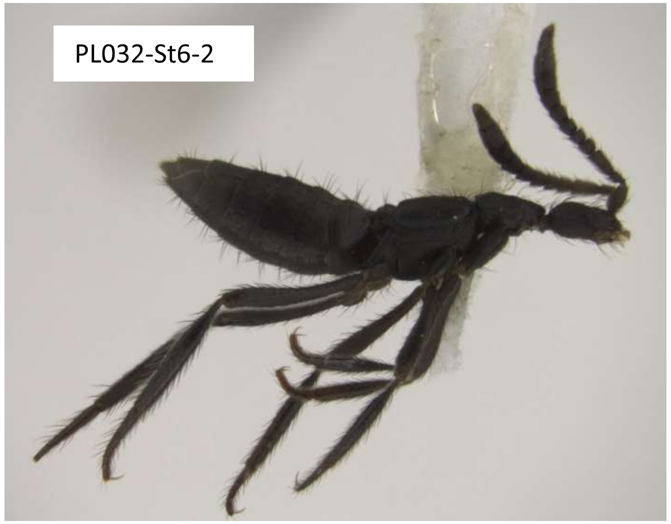

| Species B |
| --- |
| 22-ST6-4 |
| 28-ST6-1 |
| 28-ST6-3 |
| 28-ST6-4 |
| 32-ST6-1 |
| 32-ST6-2 |
| 32-ST6-3 |
| 32-ST6-5 |
| 41-ST6-2 |
| 41-ST6-3 |

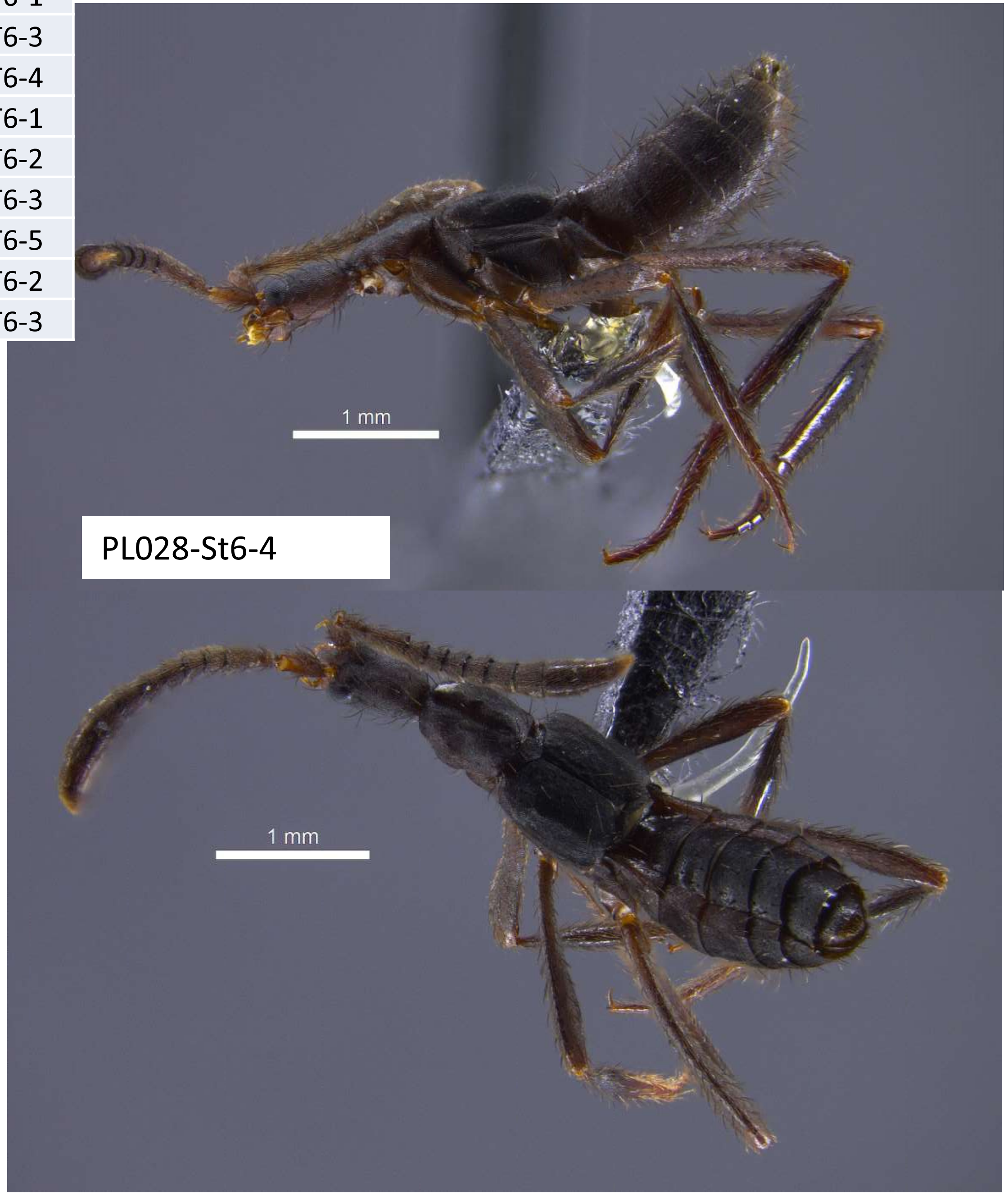


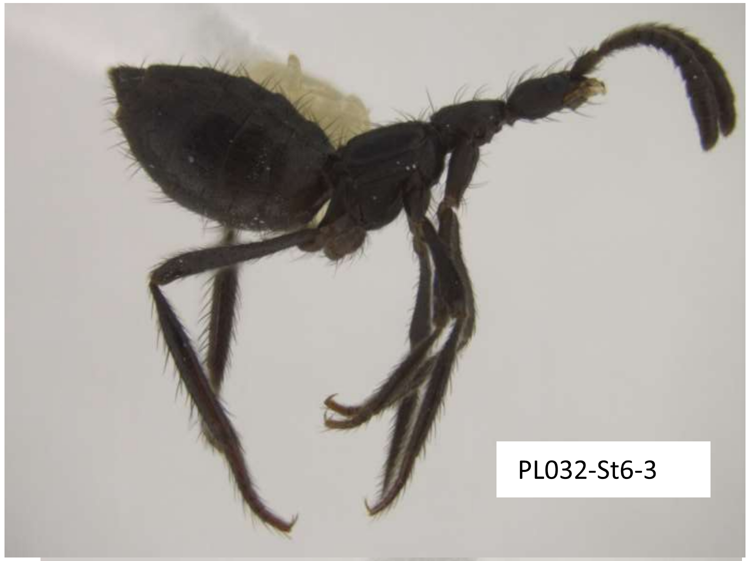


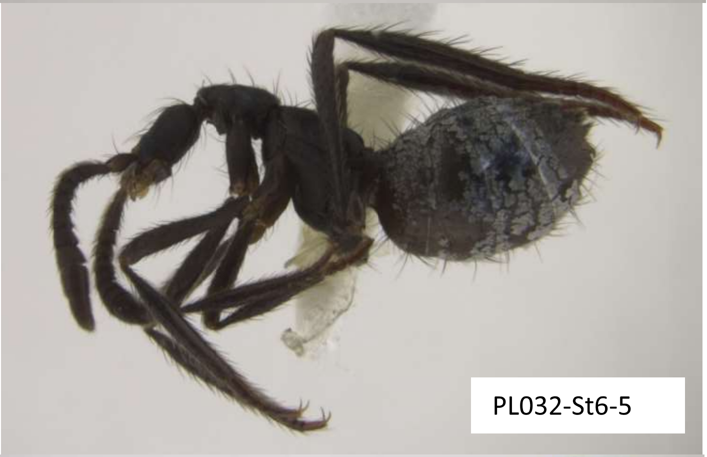


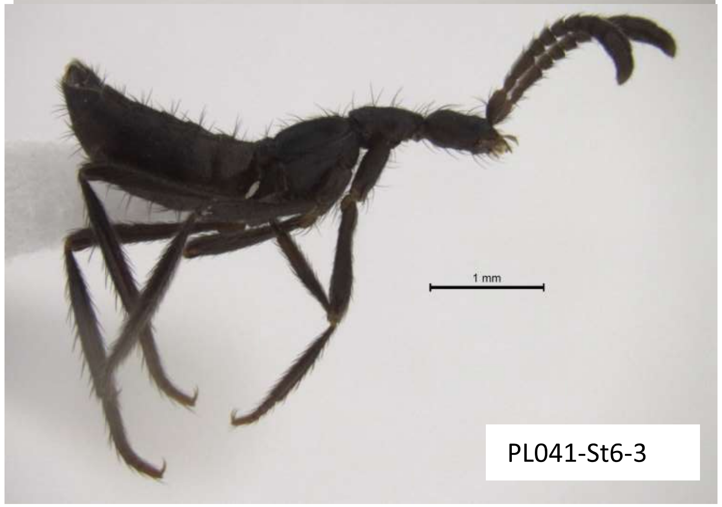

**Species ST-A? B?**

PL041-St6-1

Barcode matches ST-D, which I think is not correct… Samples confounded?

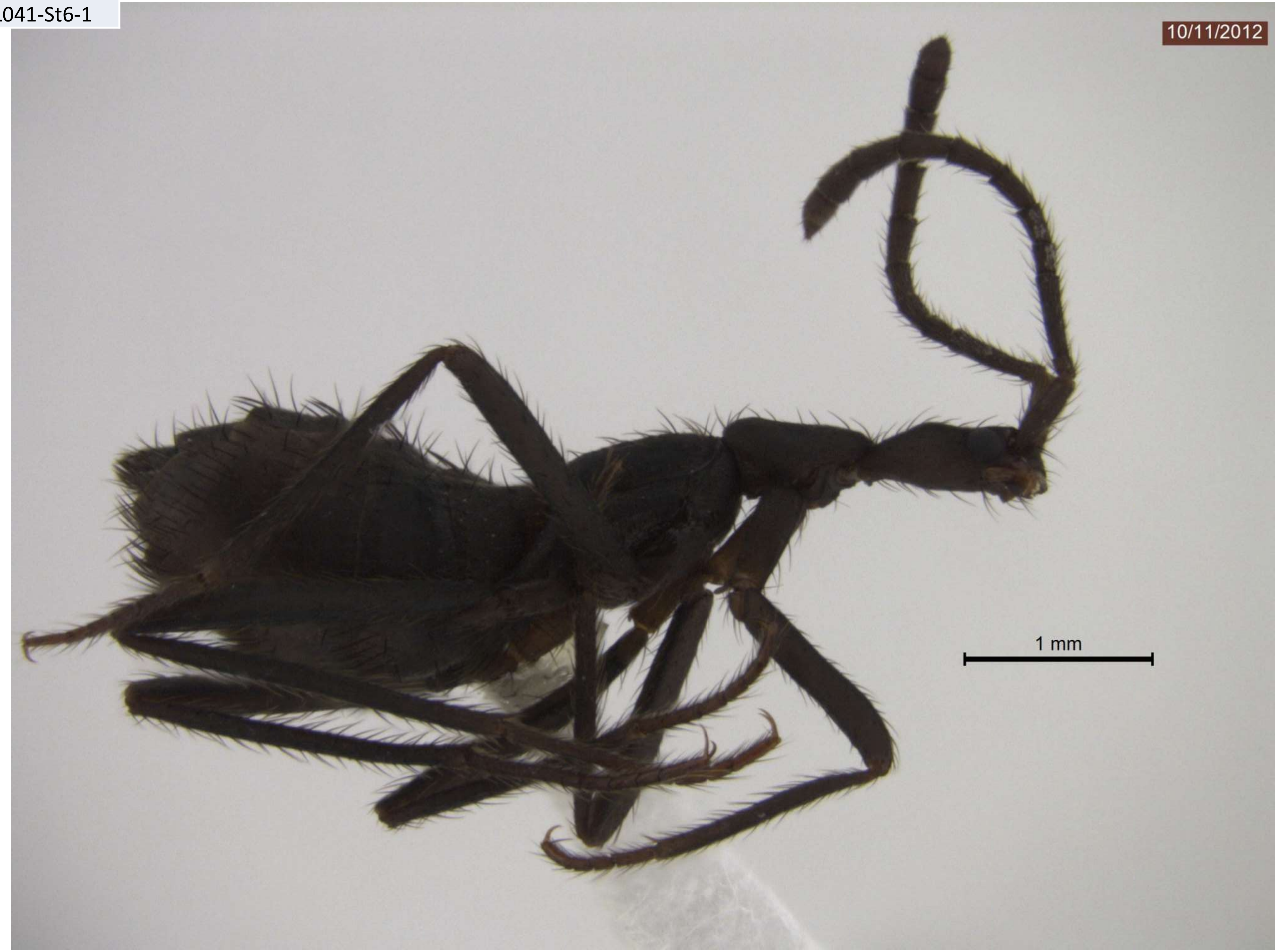

| Species ST-C |
| --- |
| PL028-ST1A-1 |
| PL028-ST1A-2 |
| PL022-ST1-1 |
| PL022-ST1-2 |
| PL028-ST1B-6 |
| PL028-ST1B-7 |
| PL028-ST1B-8 |
| PL028-ST1B-9 |
| PL028-ST1B-10 |
| PL028-ST1-S1 |
| PL028-ST1-S2 |
| PL028-ST1-S3 |
| PL028-ST1-S4 |
| PL028-ST1-S5 |
| SOD013-STA-3 |

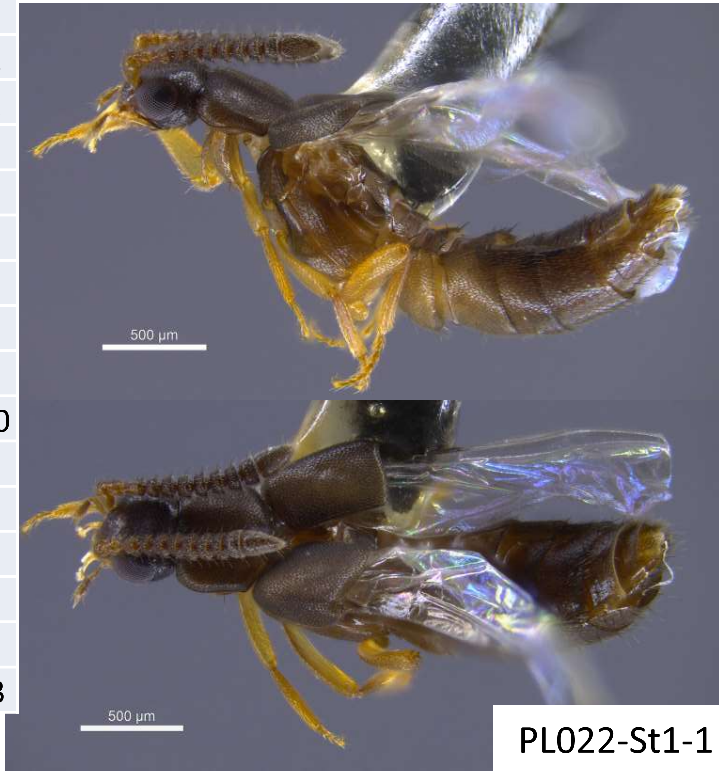


PL022-St1-1

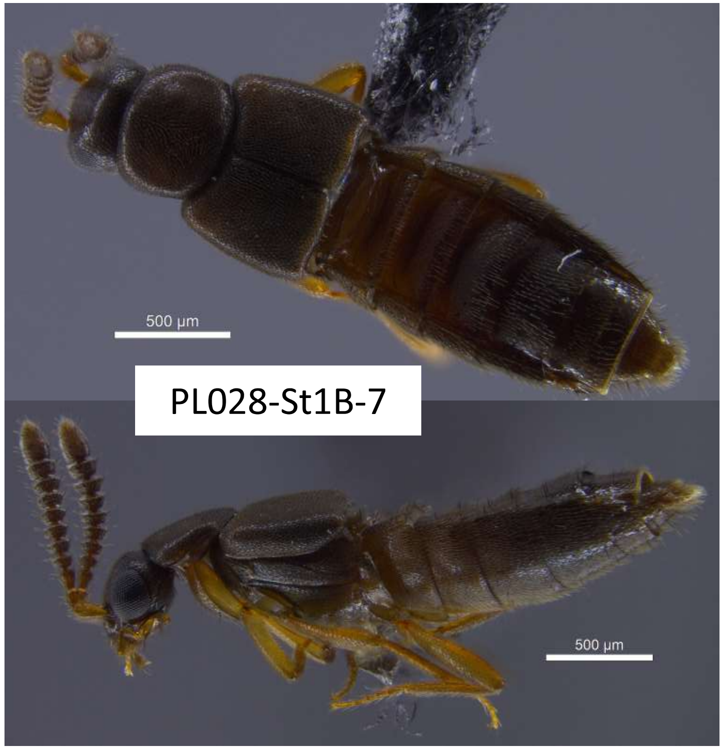


PL028-St1B-7

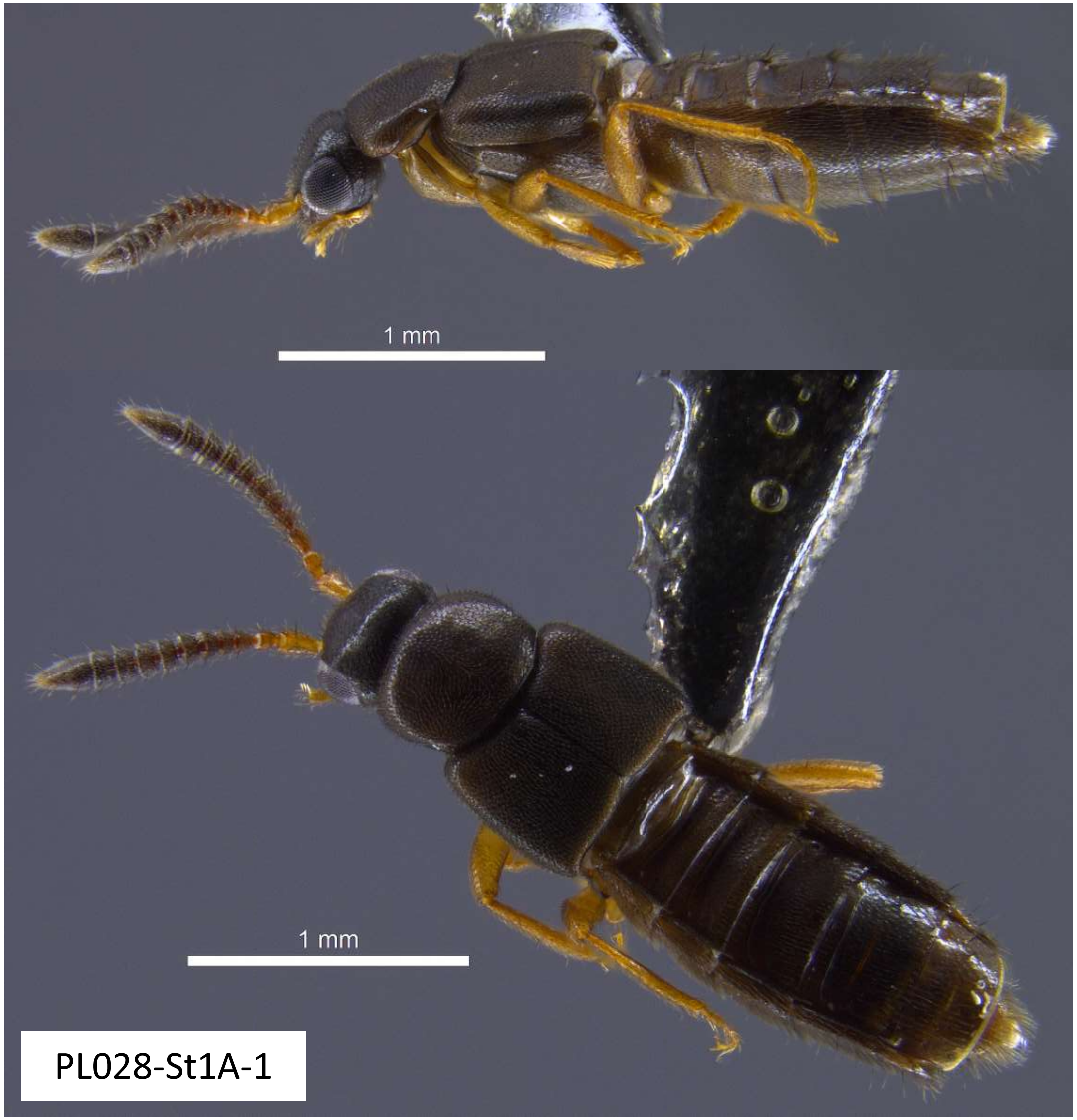


PL028-St1A-1

| Species ST-D |
| --- |
| PL022-ST5-1 |
| PL022-ST5-2 |
| PL028-ST5-1 |
| PL028-ST5-2 |

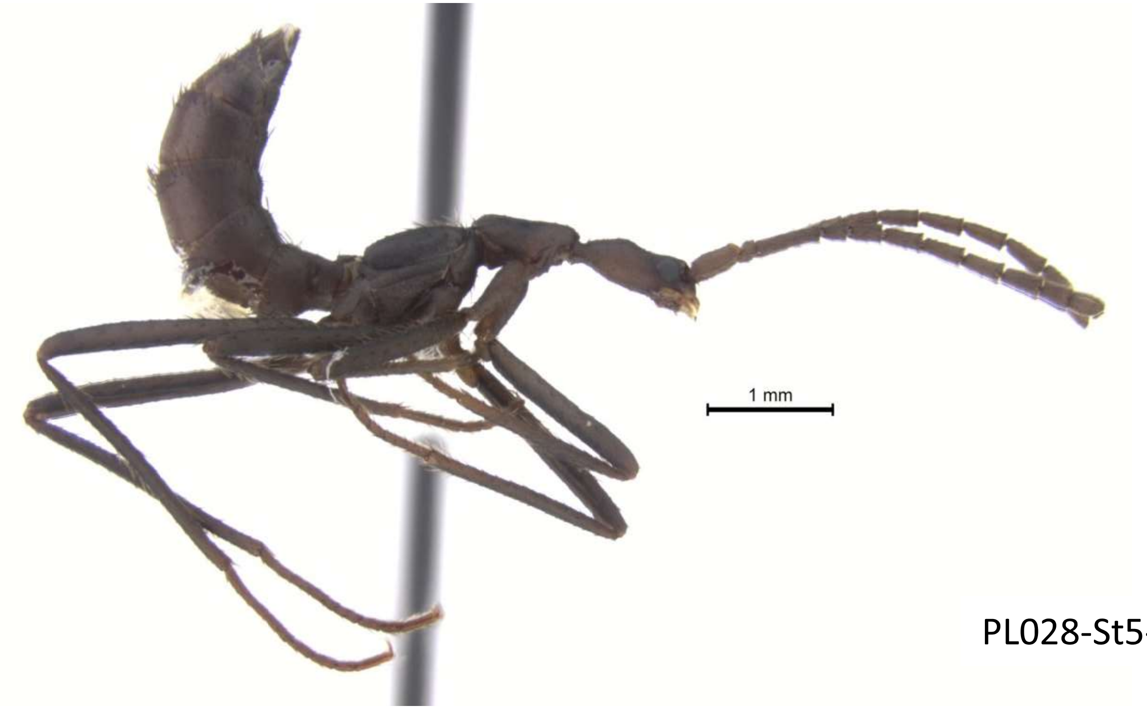


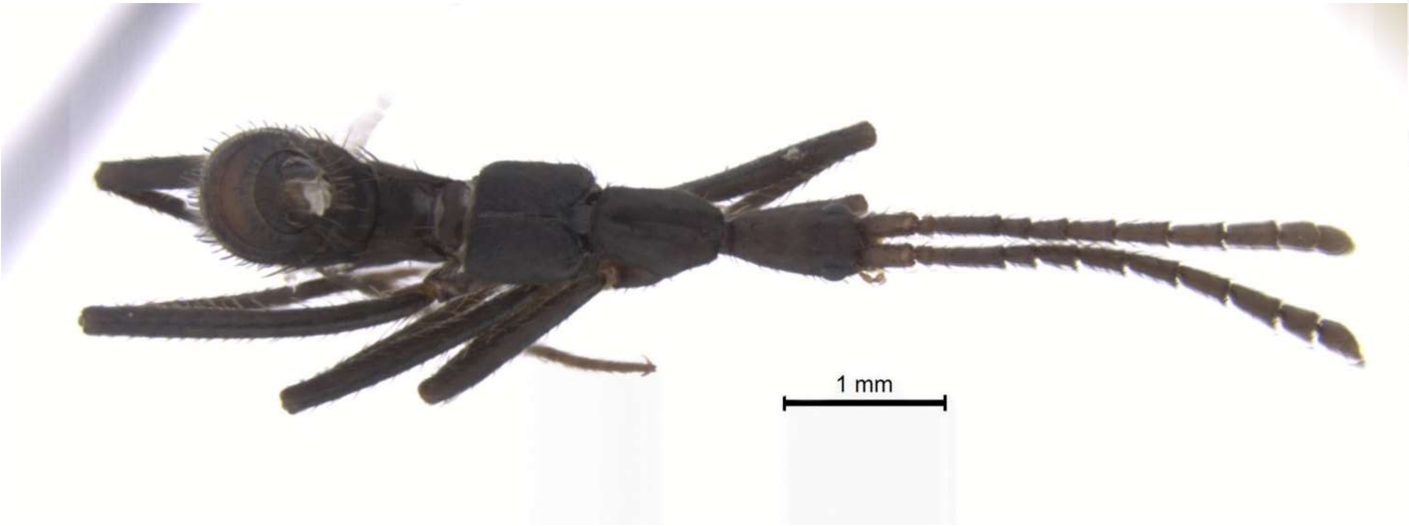


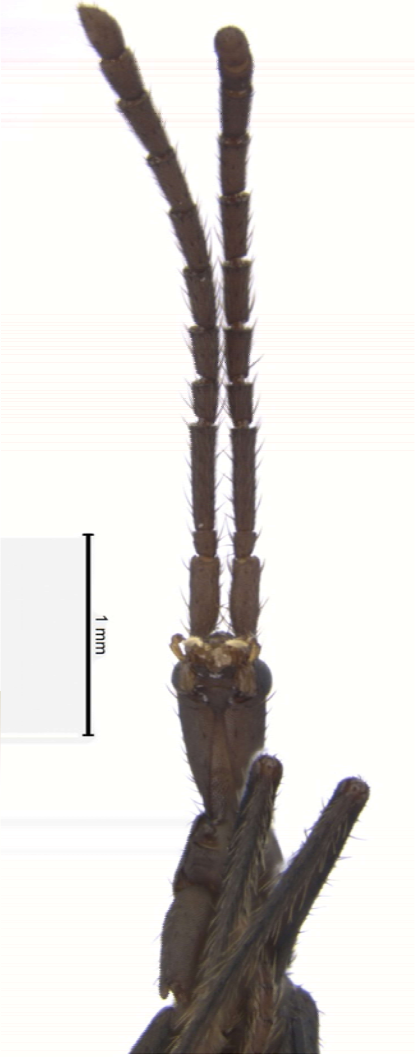


PL028-St5-1

| Species ST-E |
| --- |
| 28-ST1C-3 |
| 28-ST1C-4 |
| 28-ST1C-5 |

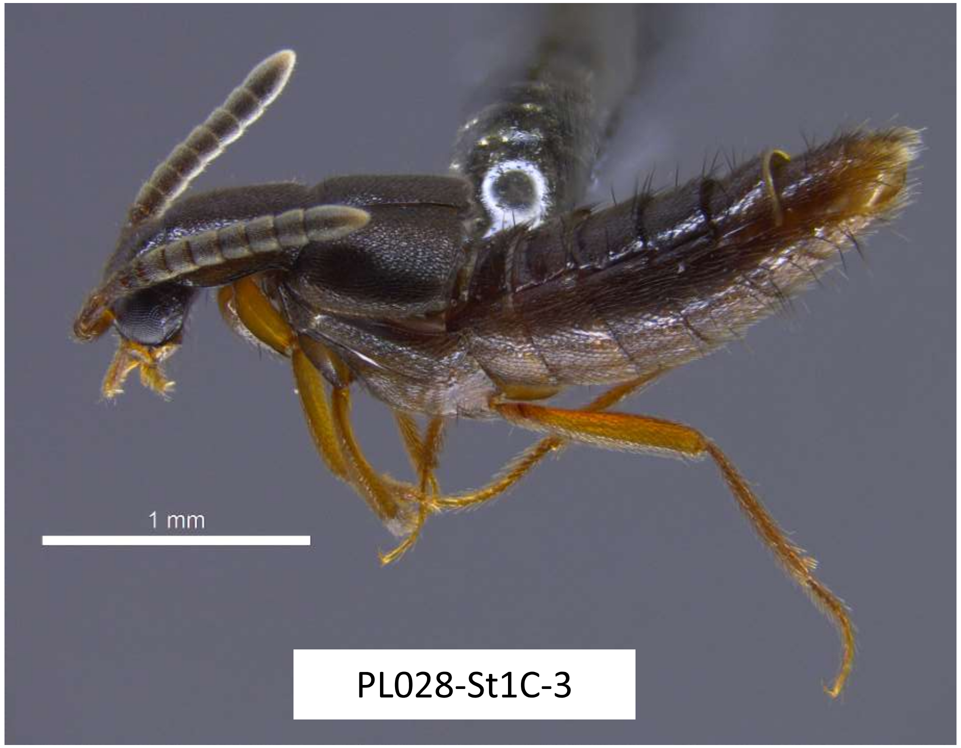


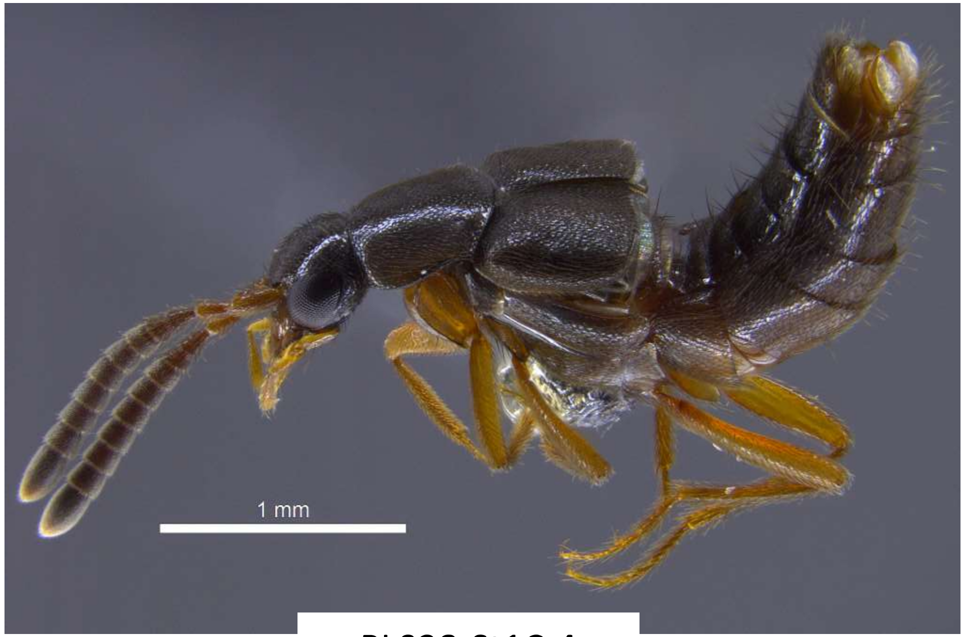


PL028-St1C-4

| Species ST-F |
| --- |
| 28-ST3E-6 |
| 28-ST3E-7 |

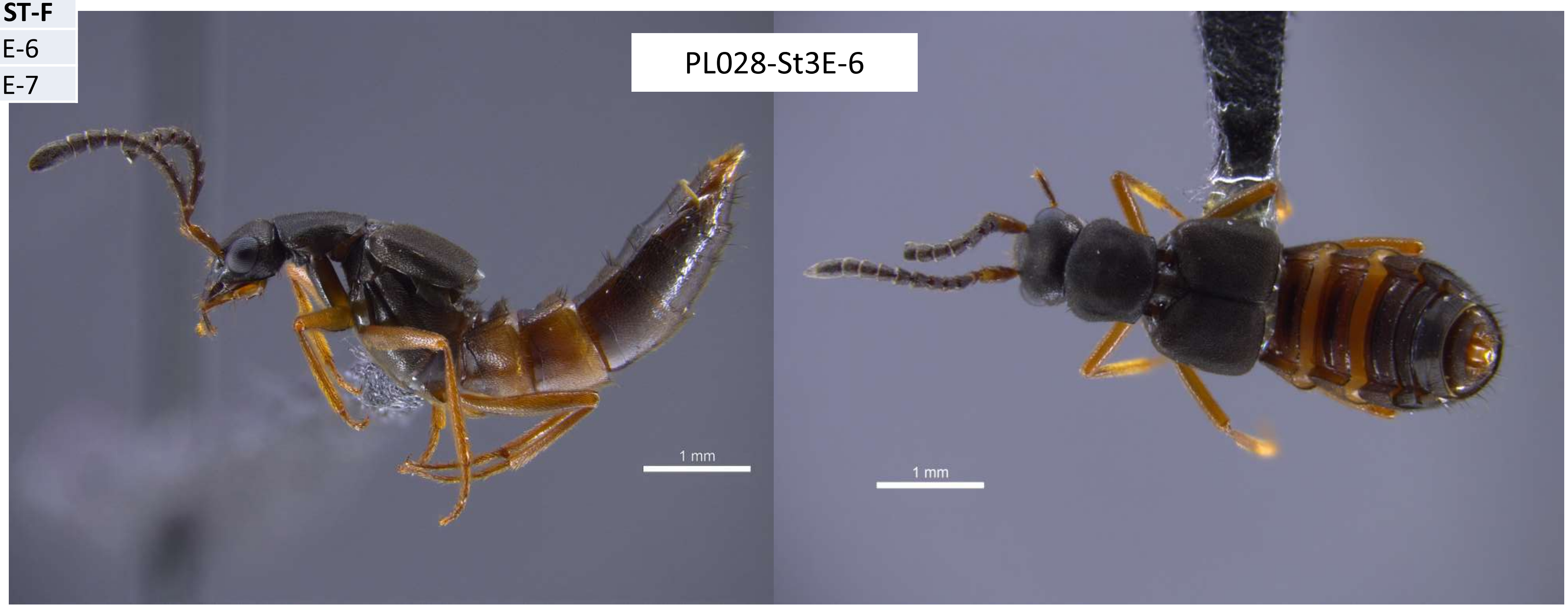


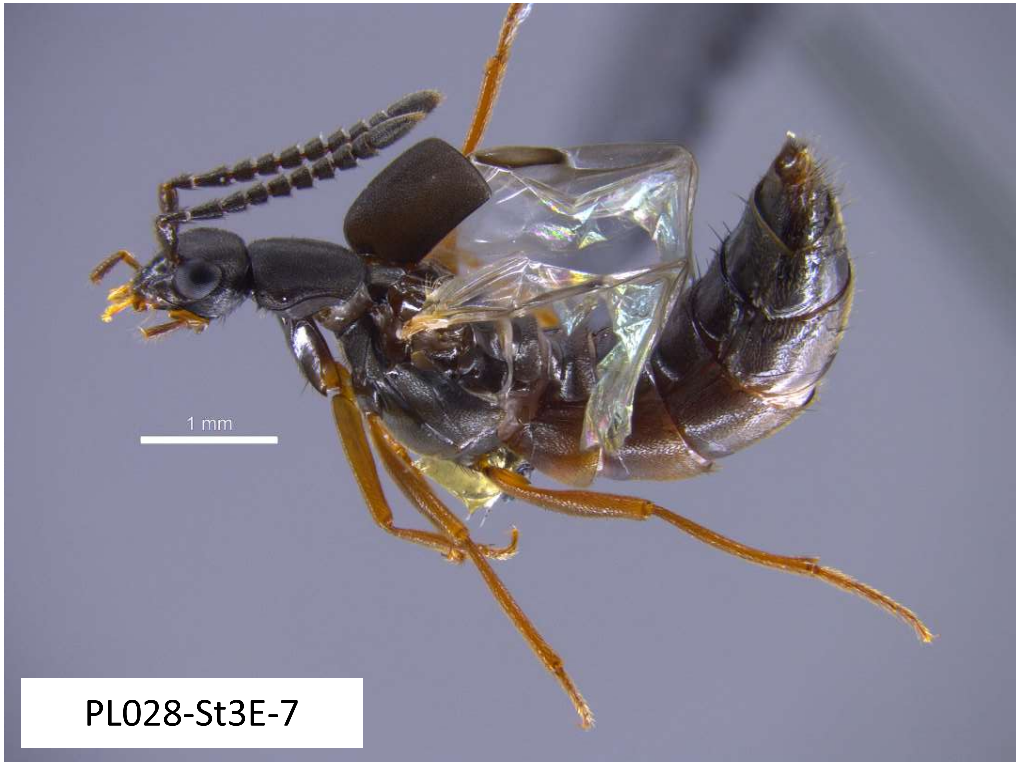

| Species ST-G |
| --- |
| PL022-ST3-1 |
| PL022-ST3-3 |
| PL022-ST3-4 |
| PL022-ST3-5 |
| PL022-ST3-6 |
| PL028-ST3-1 |
| PL028-ST3-10 |
| PL028-ST3-2 |
| PL028-ST3-3 |

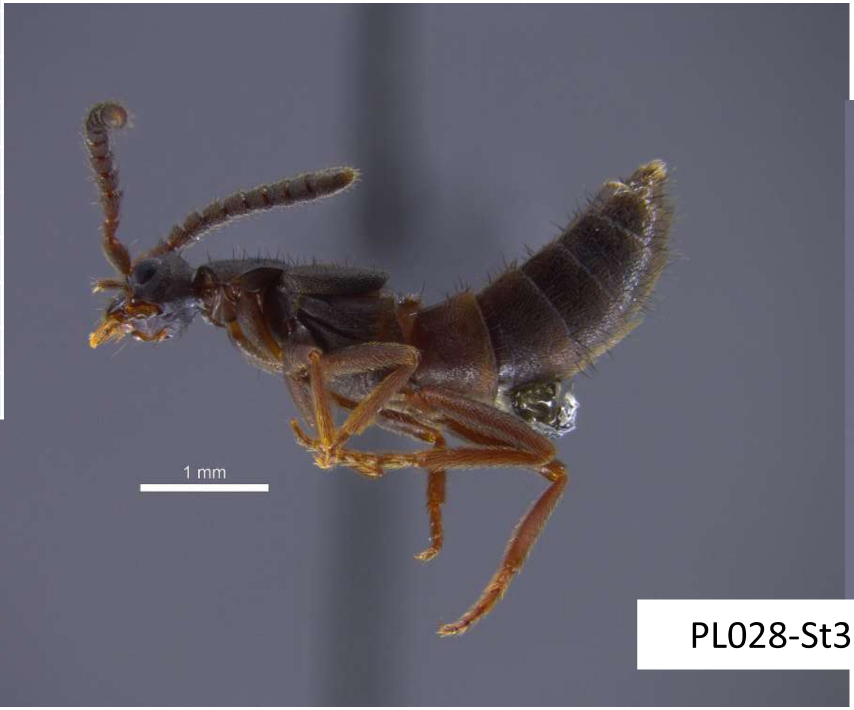


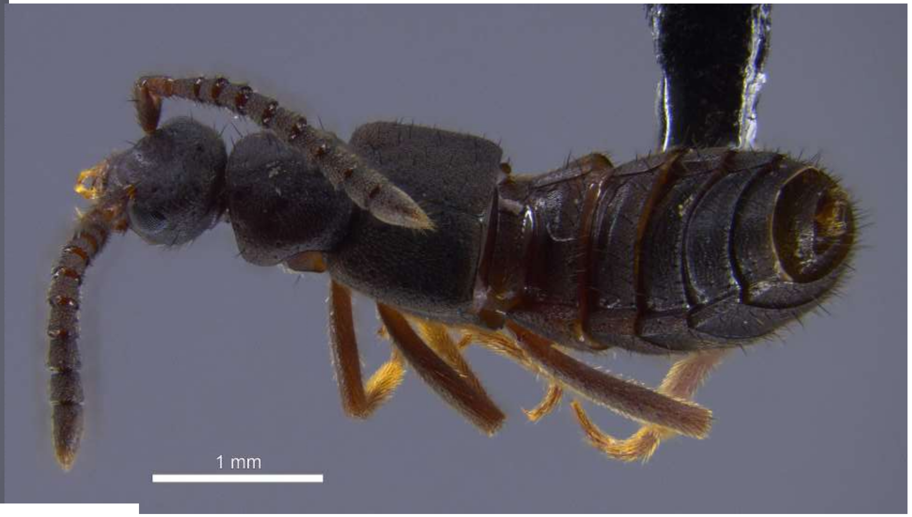


PL028-St3B-10

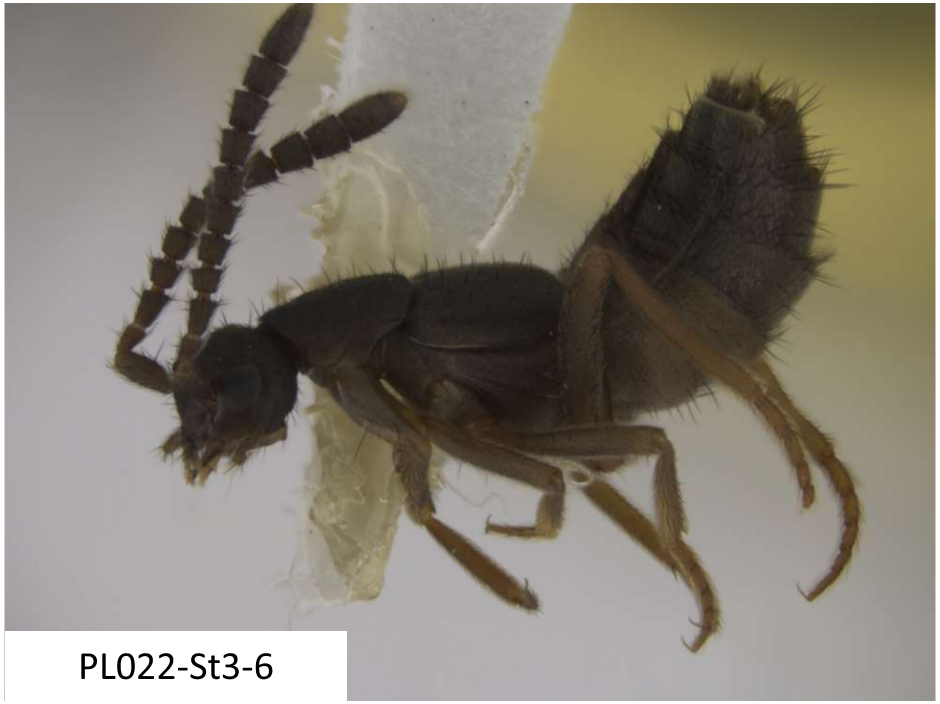

PL022-St3-6

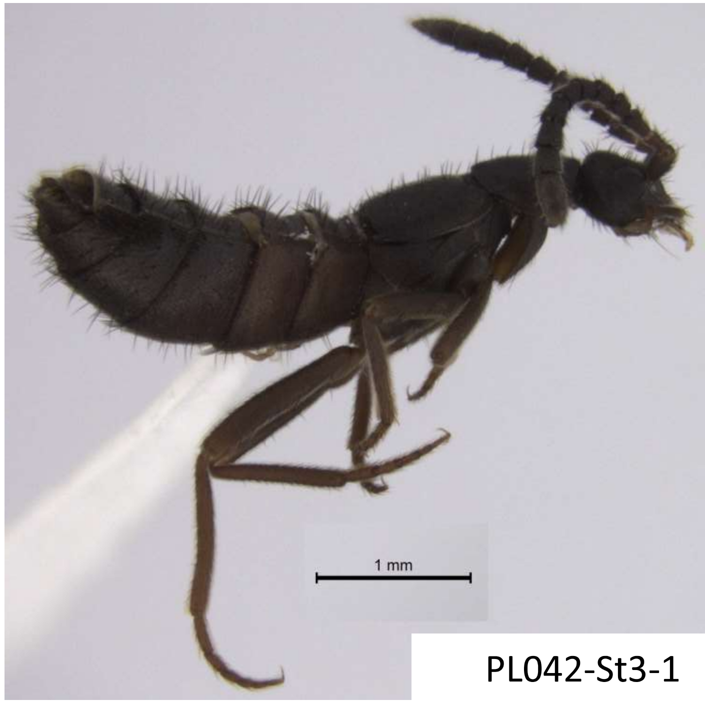


PL042-St3-1

| Species ST-G??? |
| --- |
| PL028-St3B-9 |

Morphologically appears very similar to ST-G, but the COI sequence does not align with others in the collection and BLAST-matches Silphidae most closely…

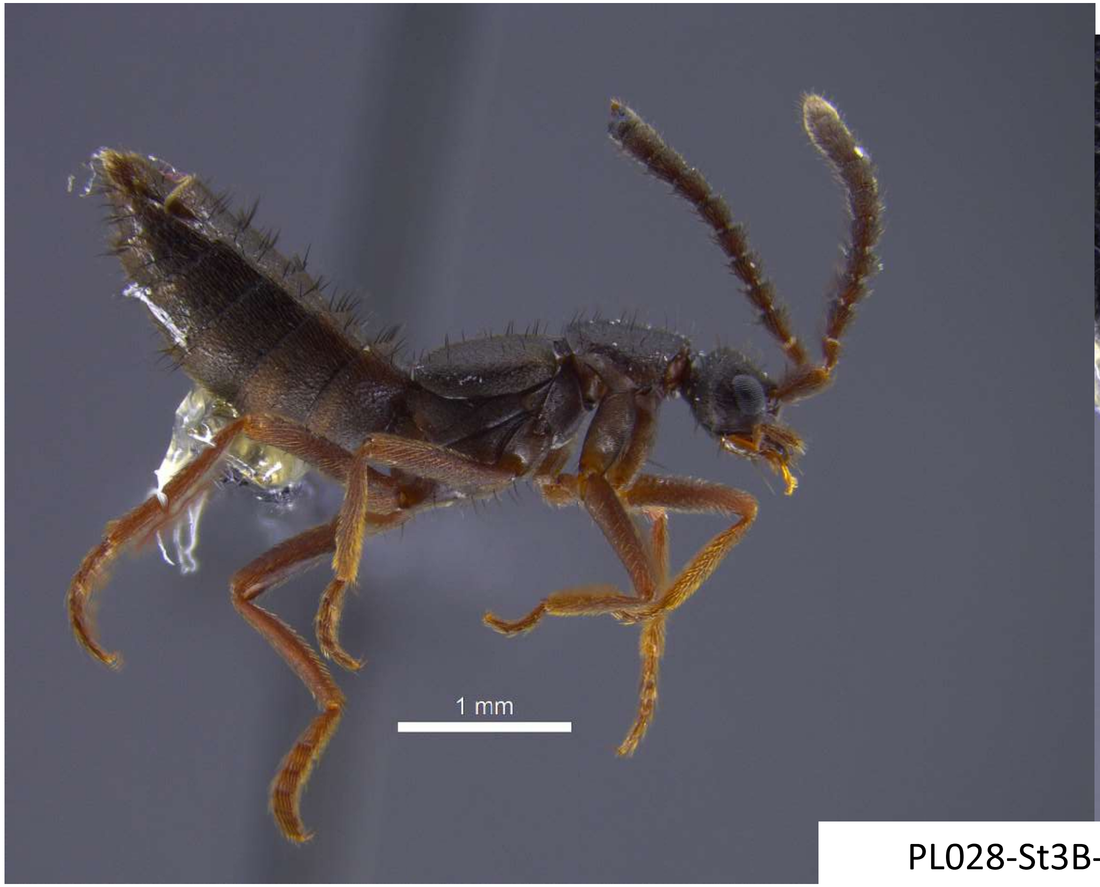


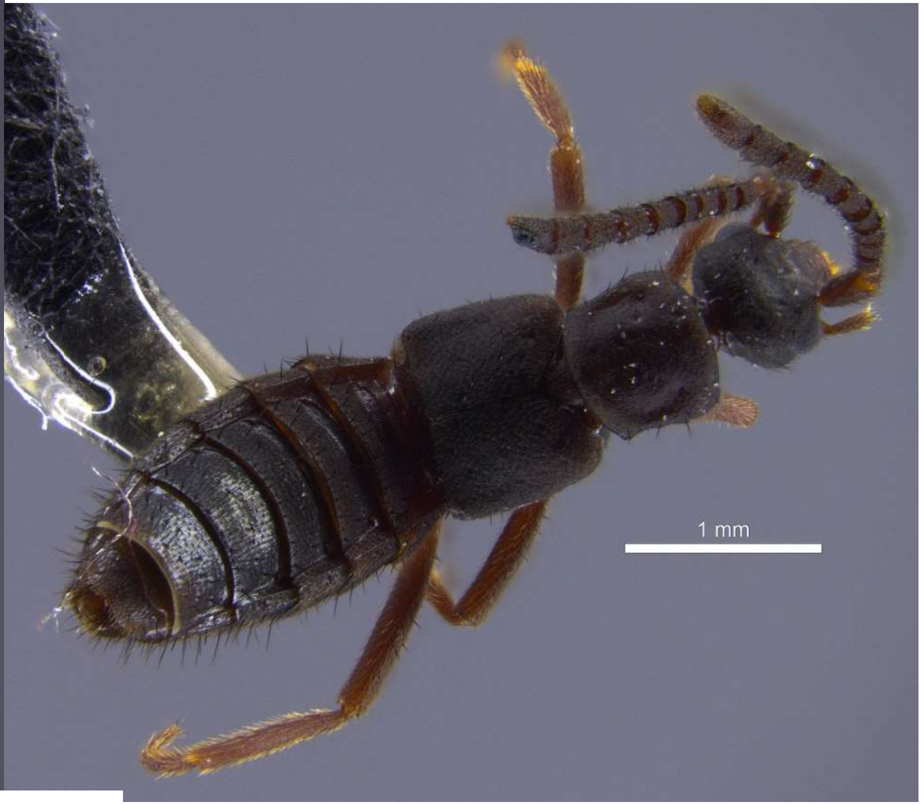


PL028-St3B-9

| Species ST-I |
| --- |
| PL022-ST2-1 |
| PL022-ST2-2 |
| PL022-ST2-3 |
| PL022-ST2-4 |
| PL028-ST2-1 |
| PL028-ST2-2 |
| PL028-ST2-3 |
| PL028-ST2-4 |
| PL028-ST2-5 |
| SOD013-STA-B |

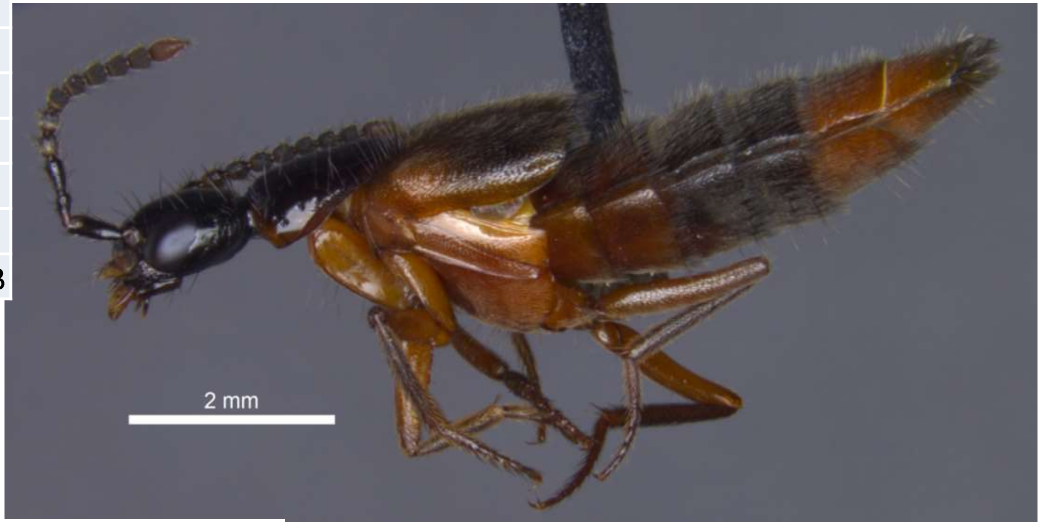


PL022-St2-1

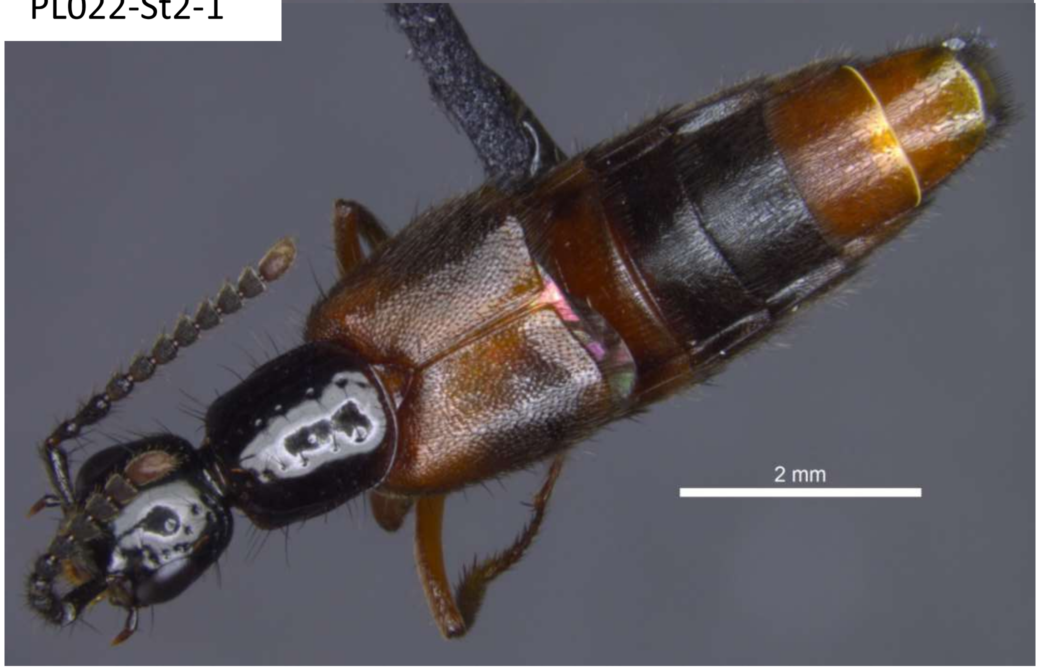


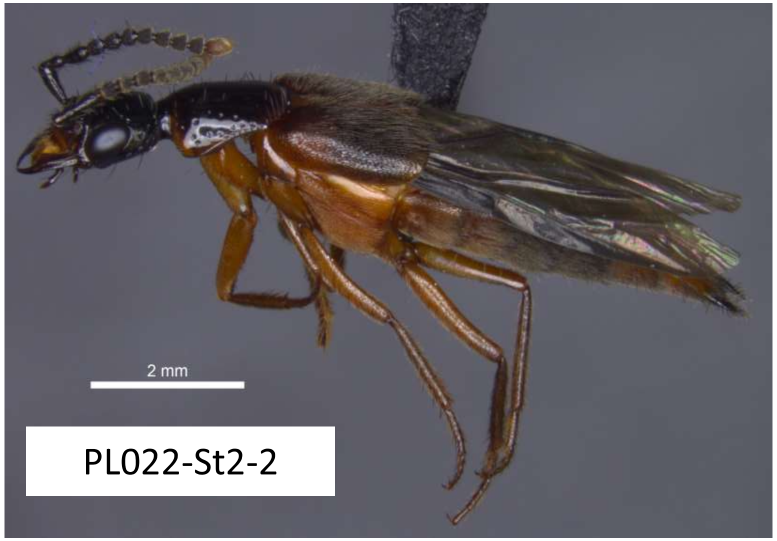


PL022-St2-2

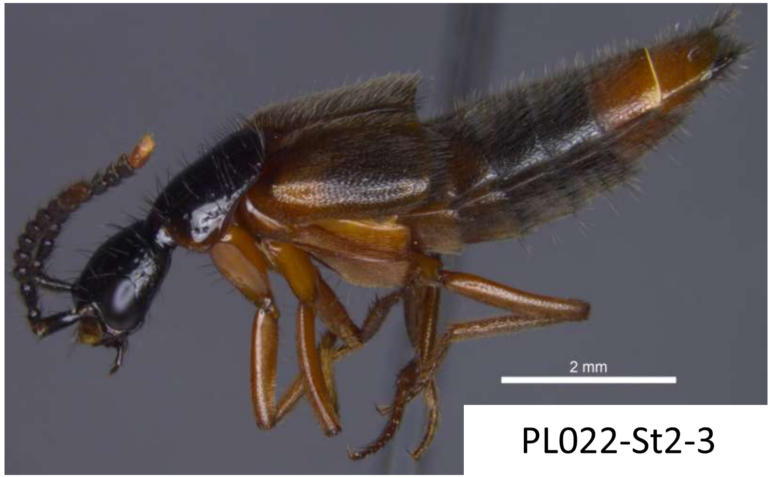


PL022-St2-3

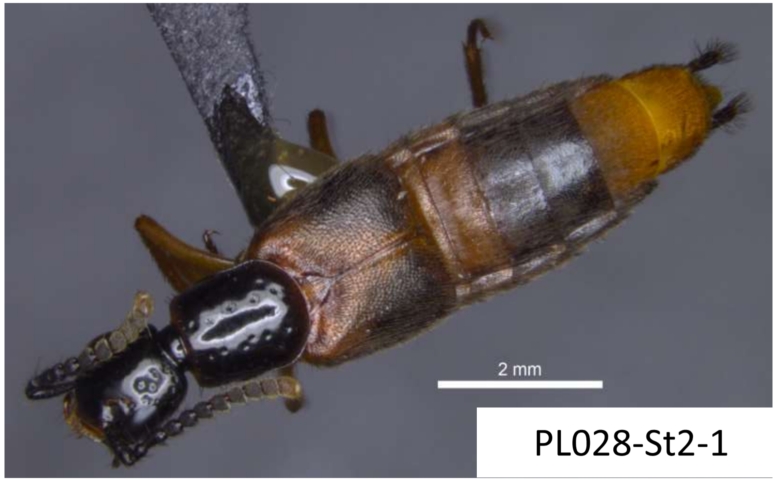


PL028-St2-1

**Species B1**

28-B1-1

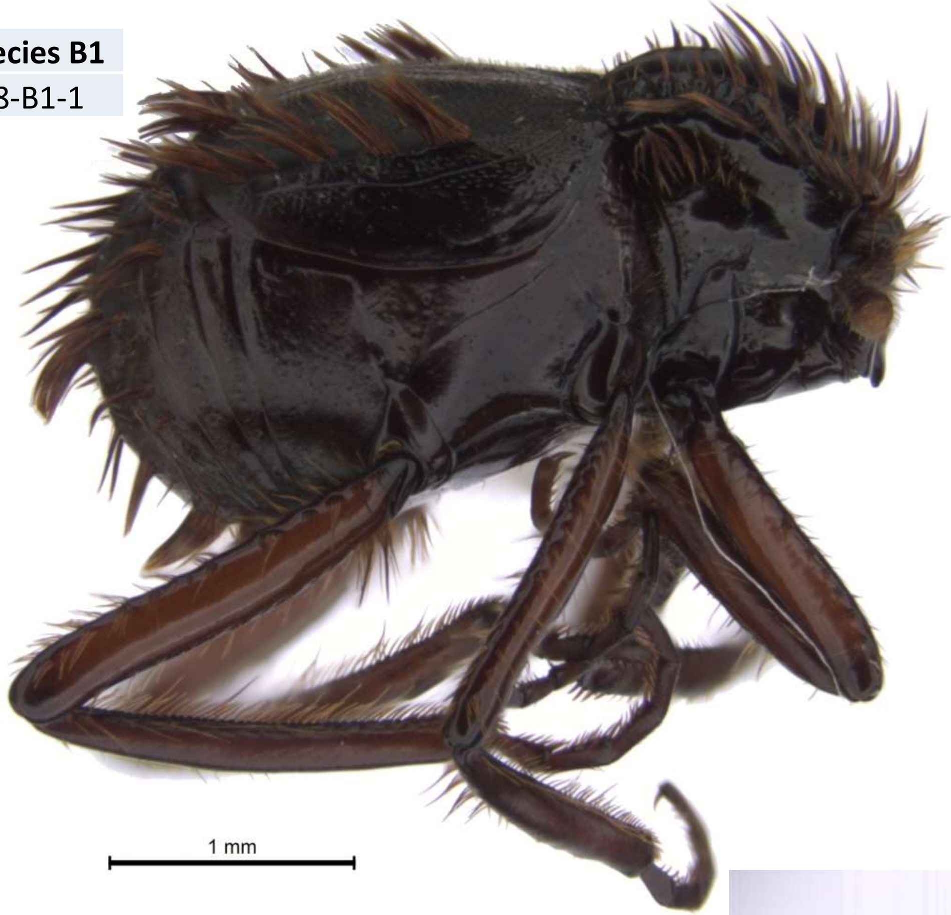


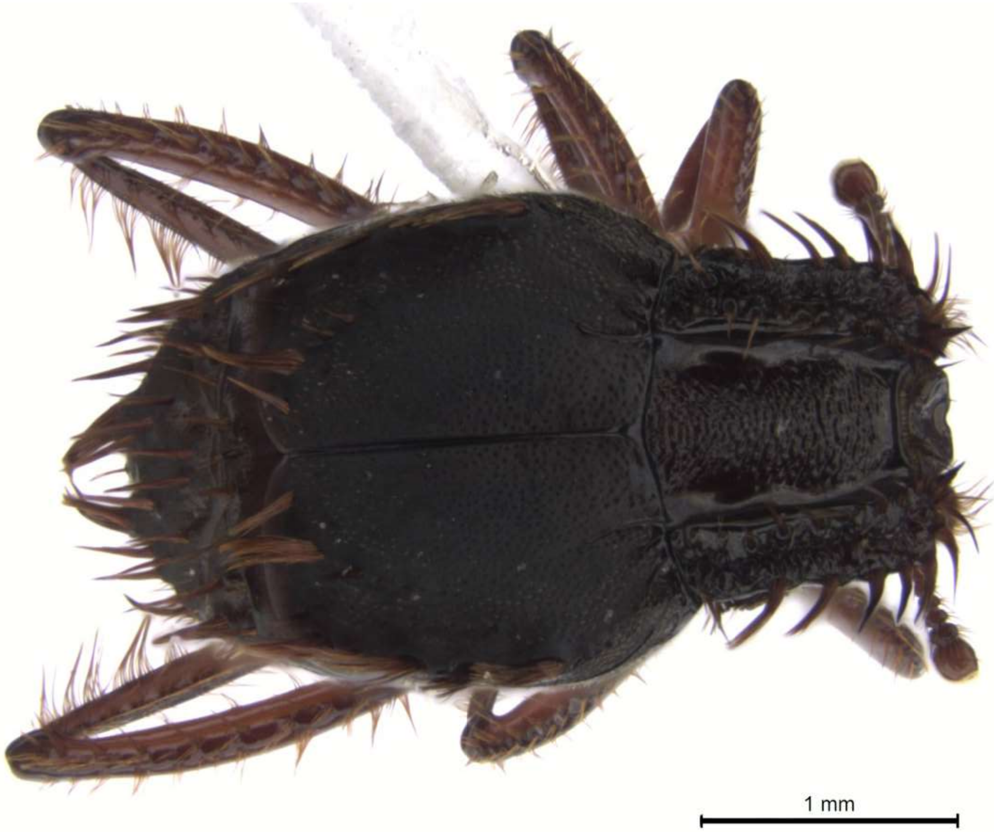


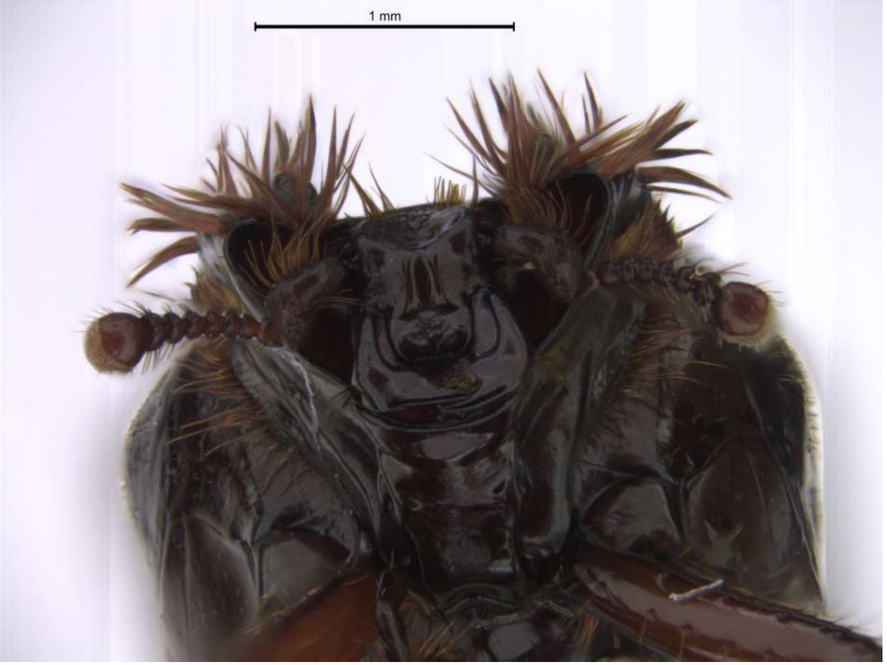

**Species B2**

28-B2-1

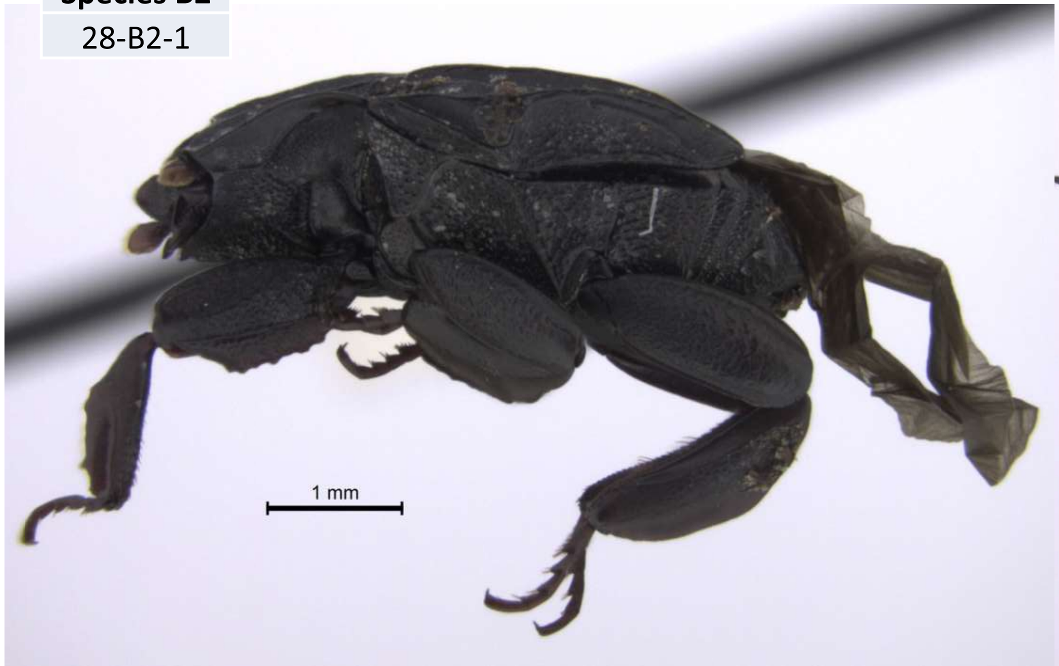


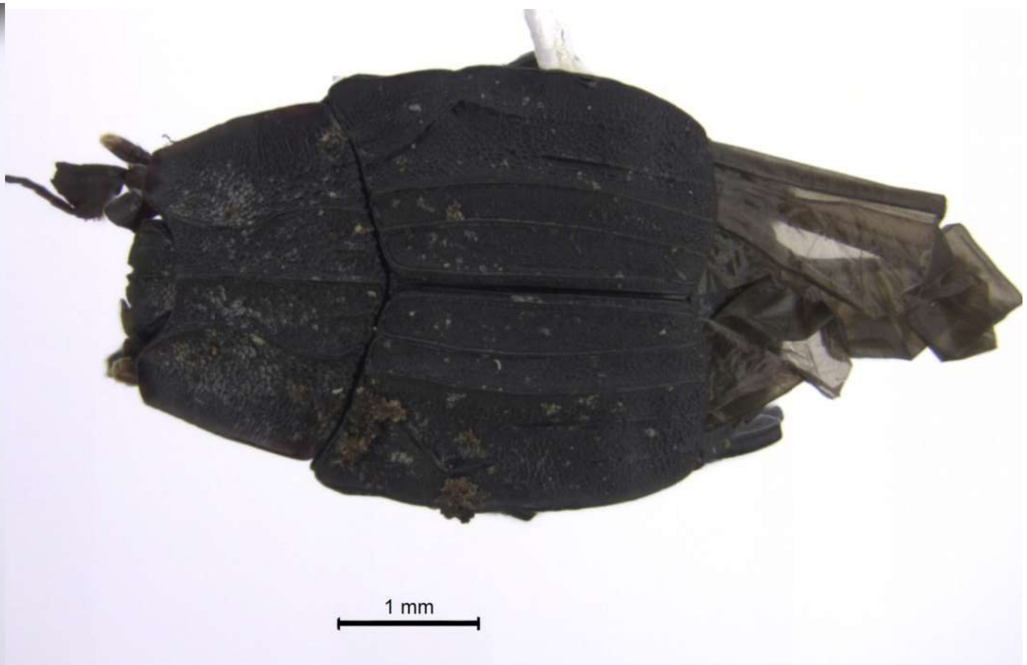


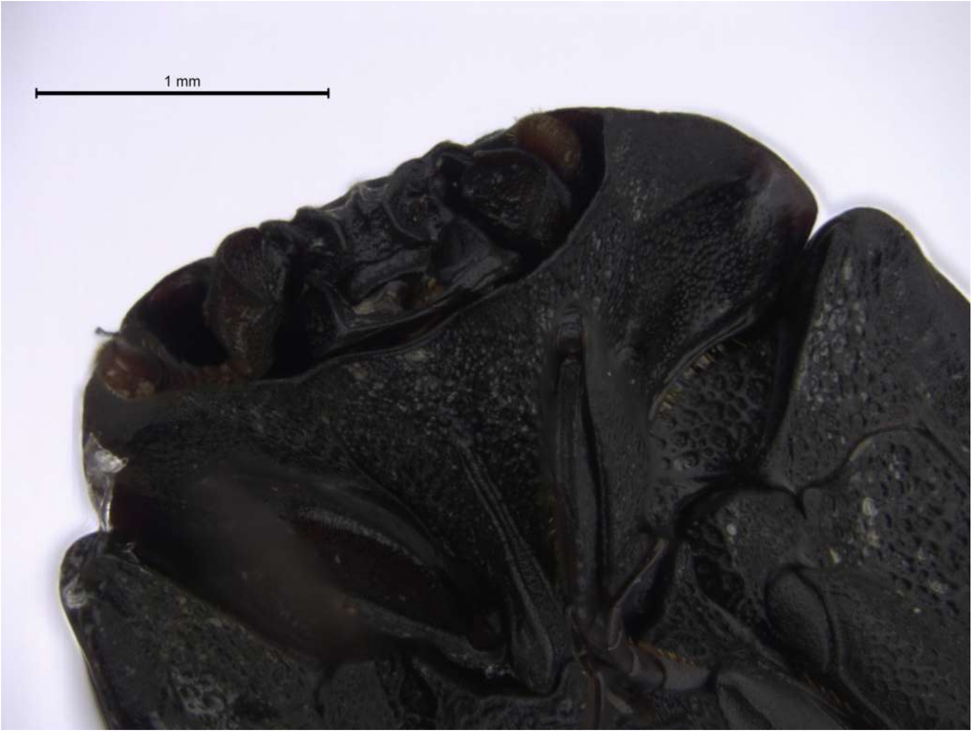


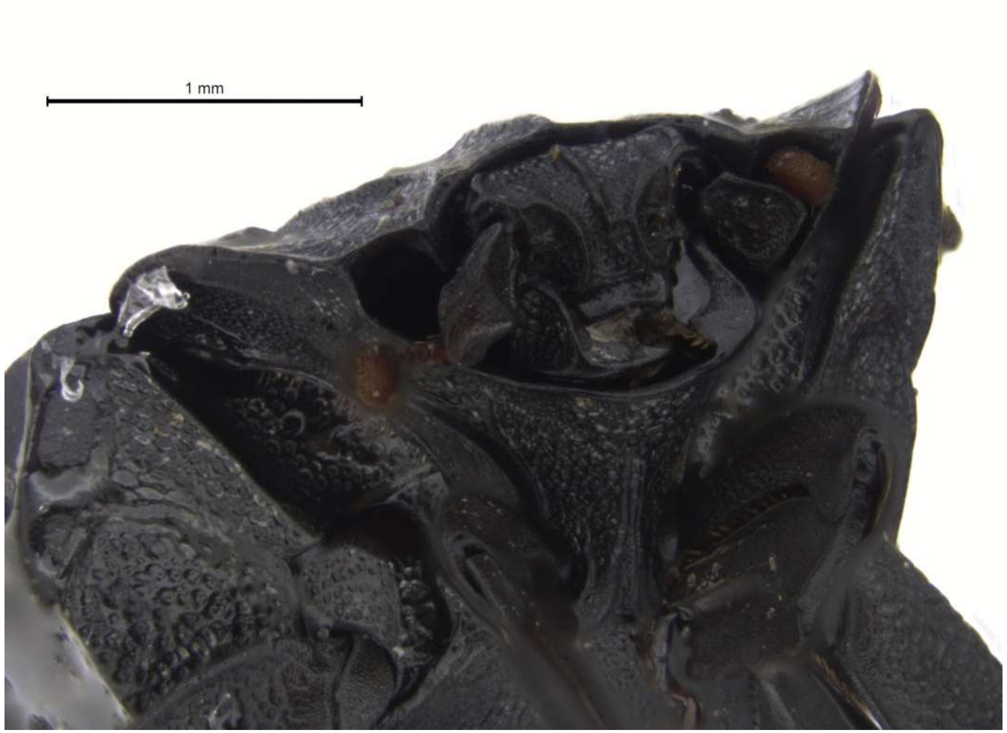

| Species B3 |
| --- |
| 22-B3-1 |

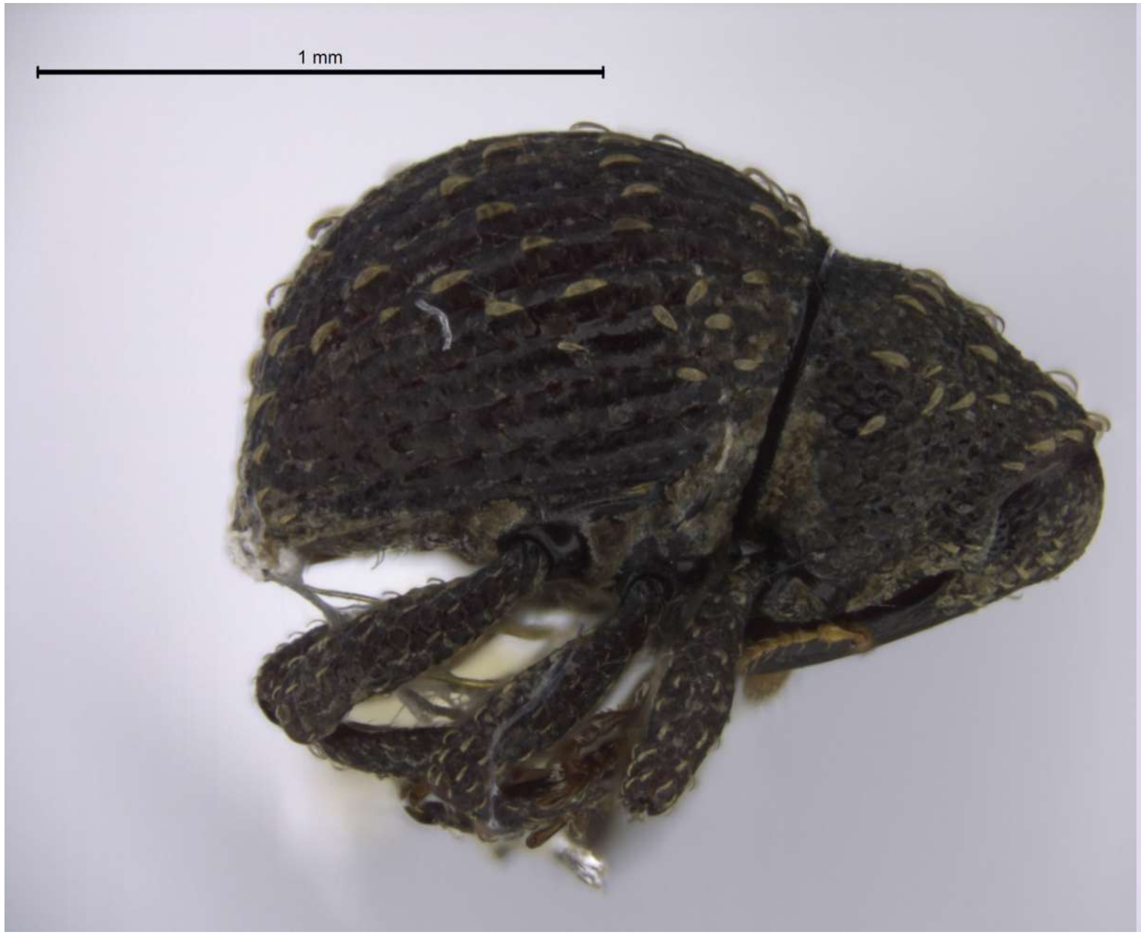


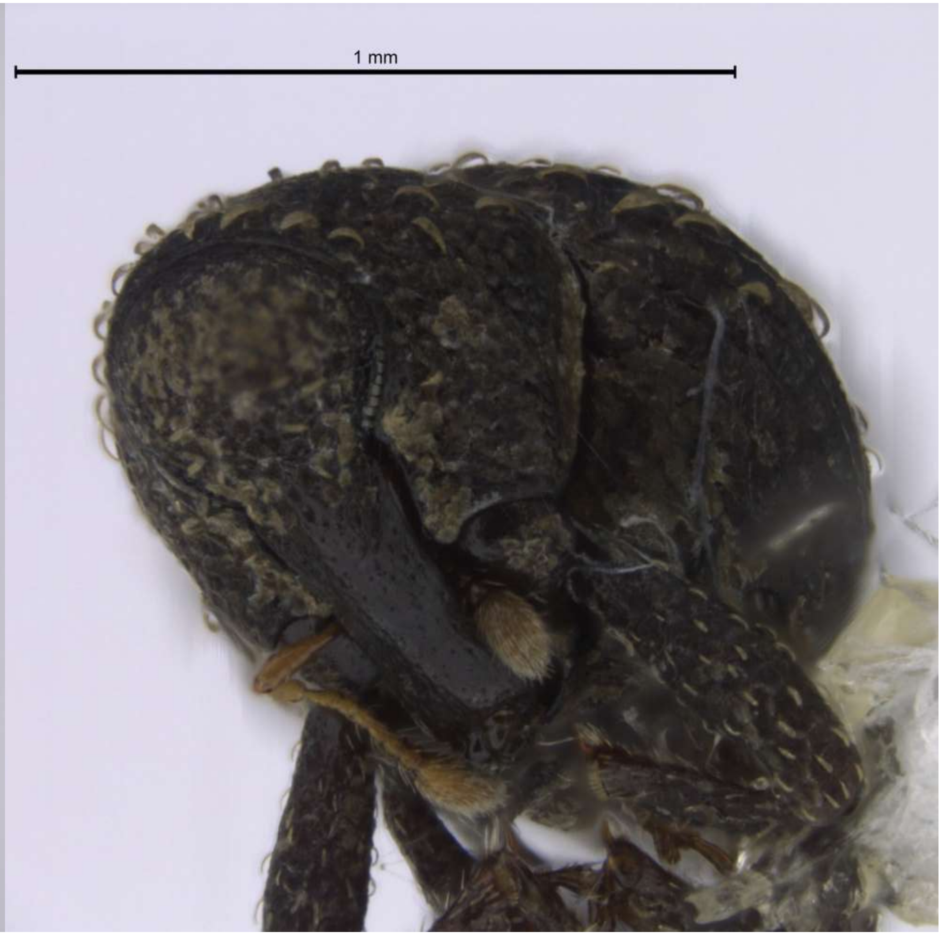

| Species HIST |
| --- |
| SOD013-HIS-1 |

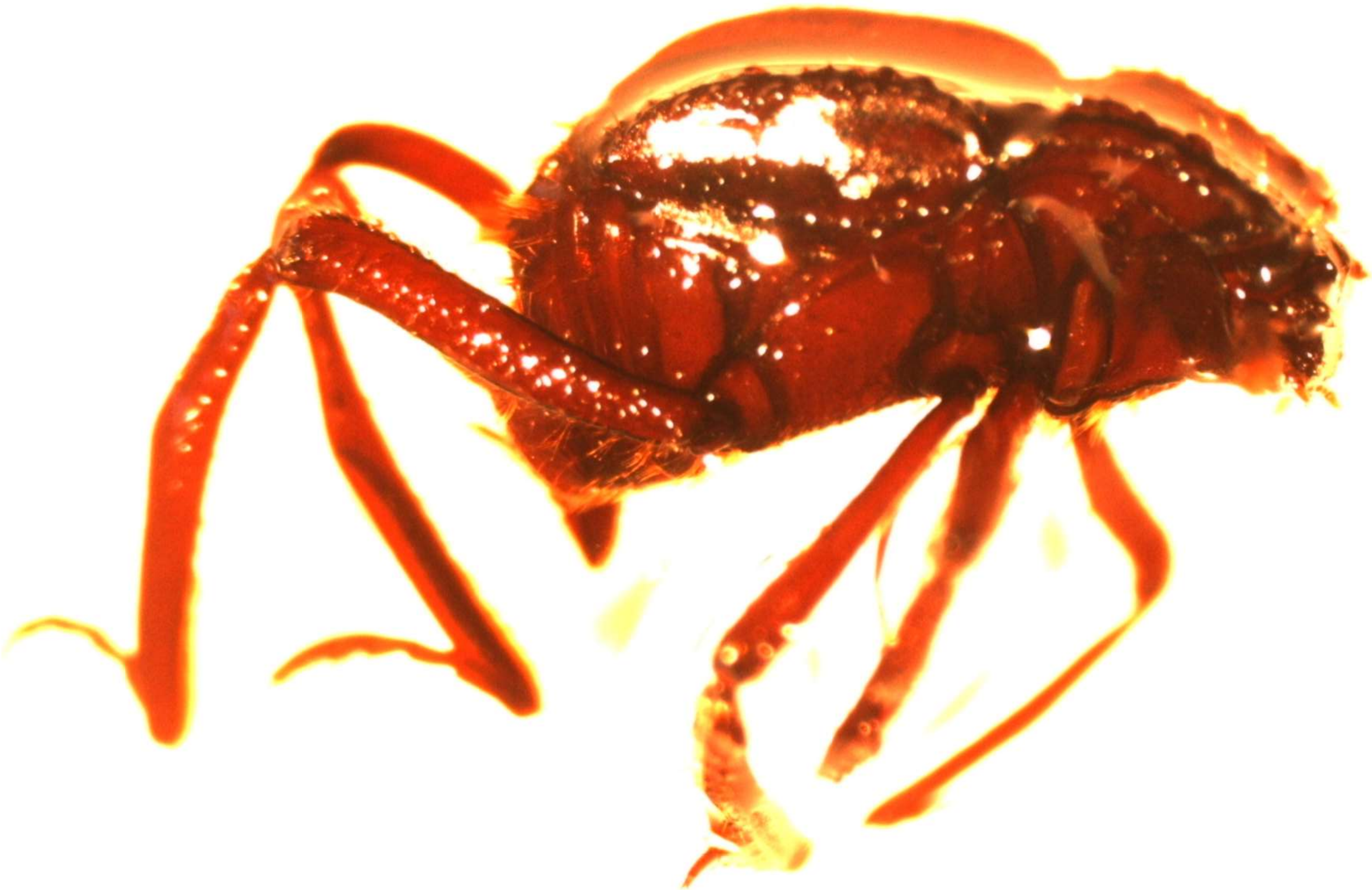

| Species FRE-12 |
| --- |
| 47-FRE-1 |
| 47-FRE-2 |

Non-myrmecophile – caught in forest litter away from army ant colonies

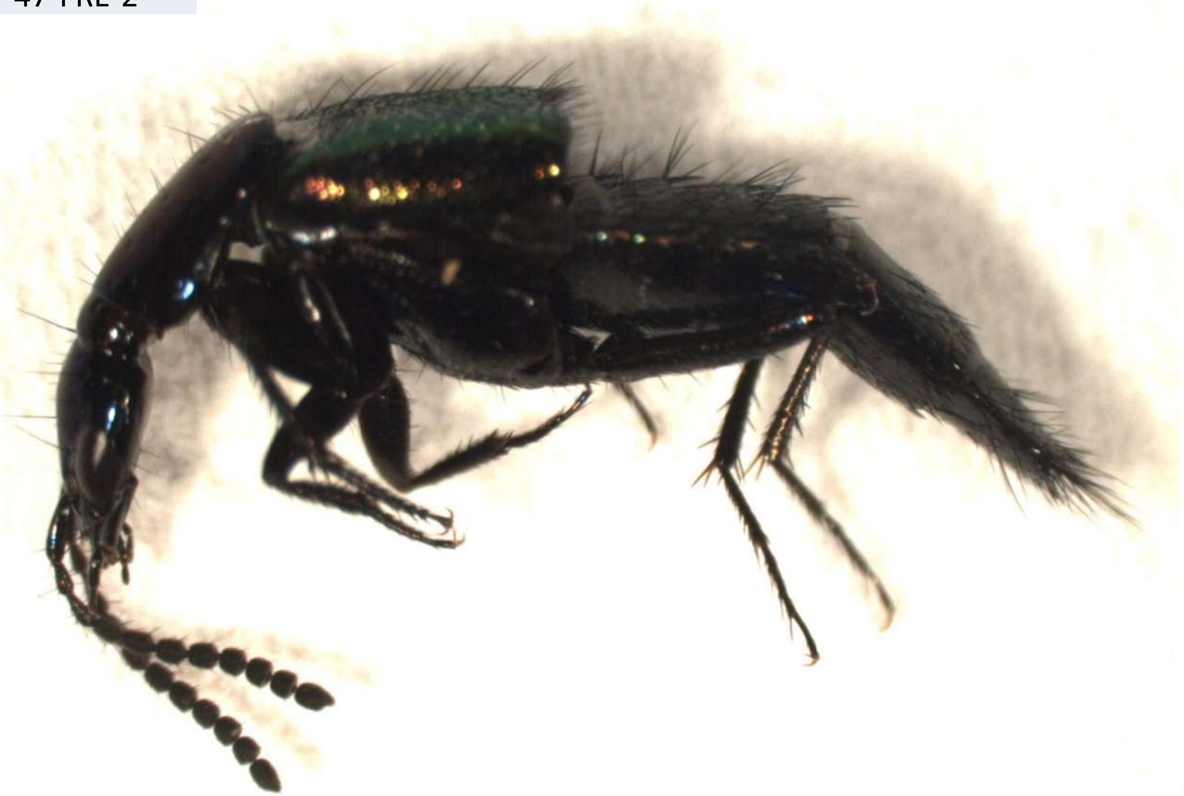

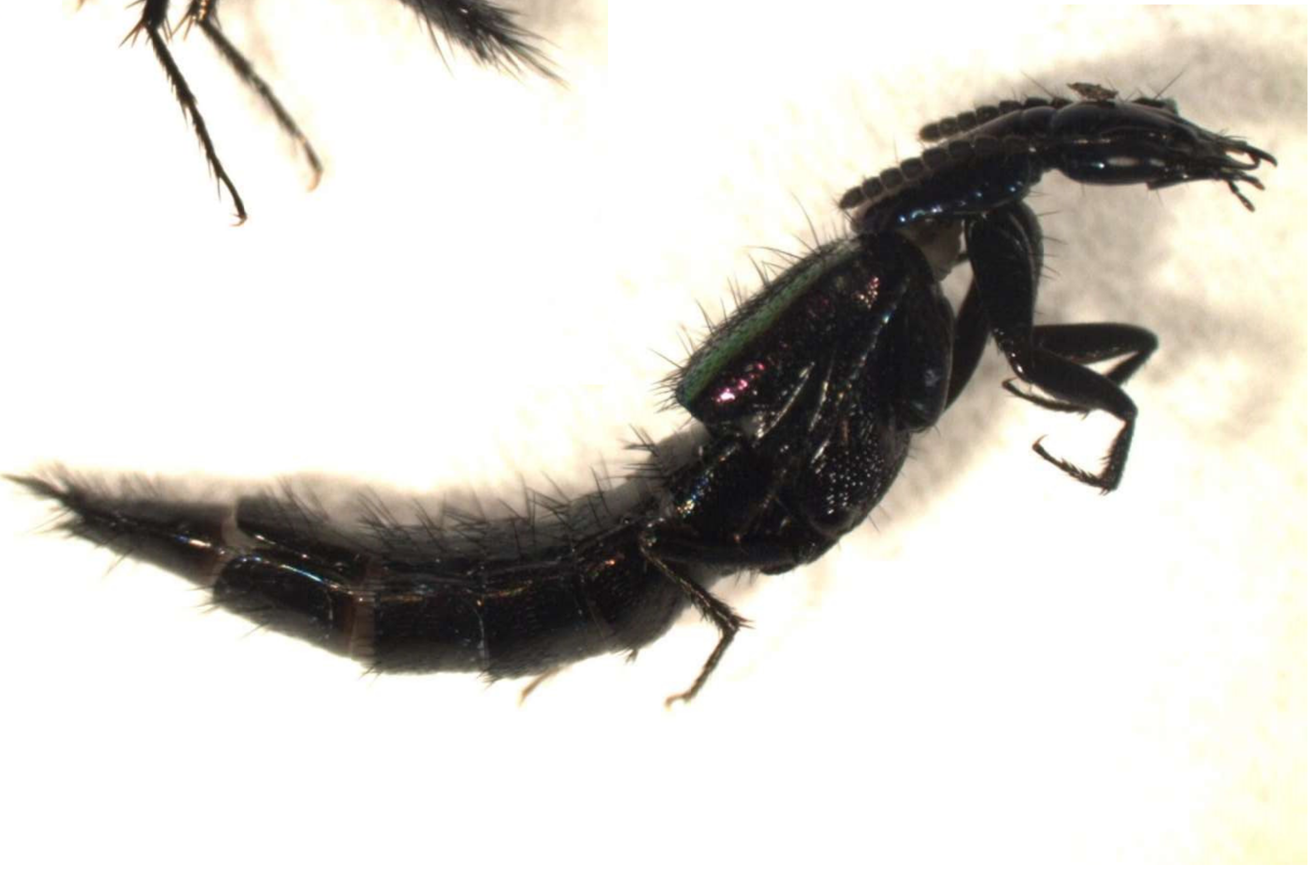

| Species FRE-34 |
| --- |
| 47-FRE-3 |
| 47-FRE-4 |

Non-myrmecophile – caught in forest litter away from army ant colonies

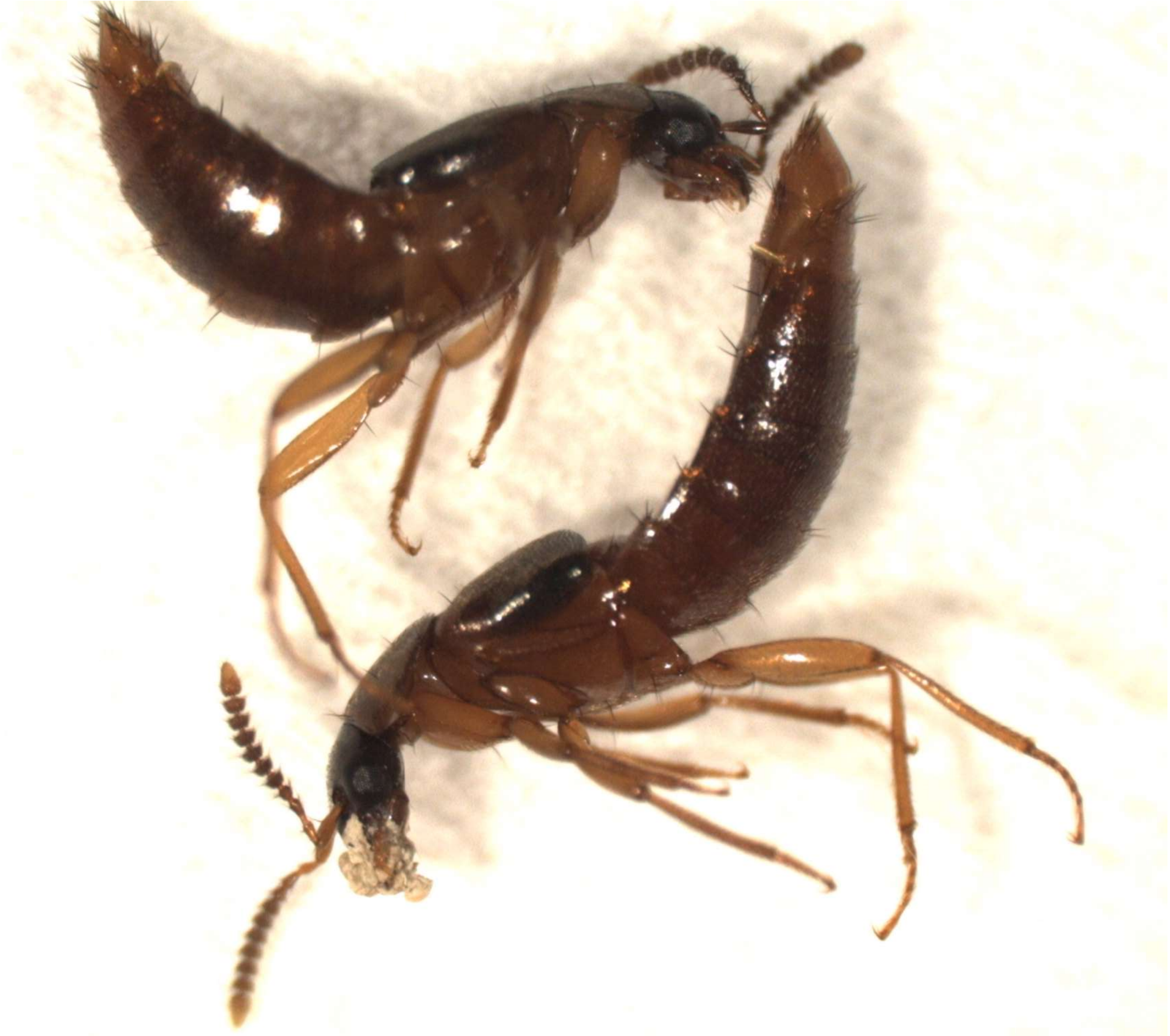

| Species FRE-5 |
| --- |
| PL047-FRE-5 |

Non-myrmecophile – caught in forest litter away from army ant colonies

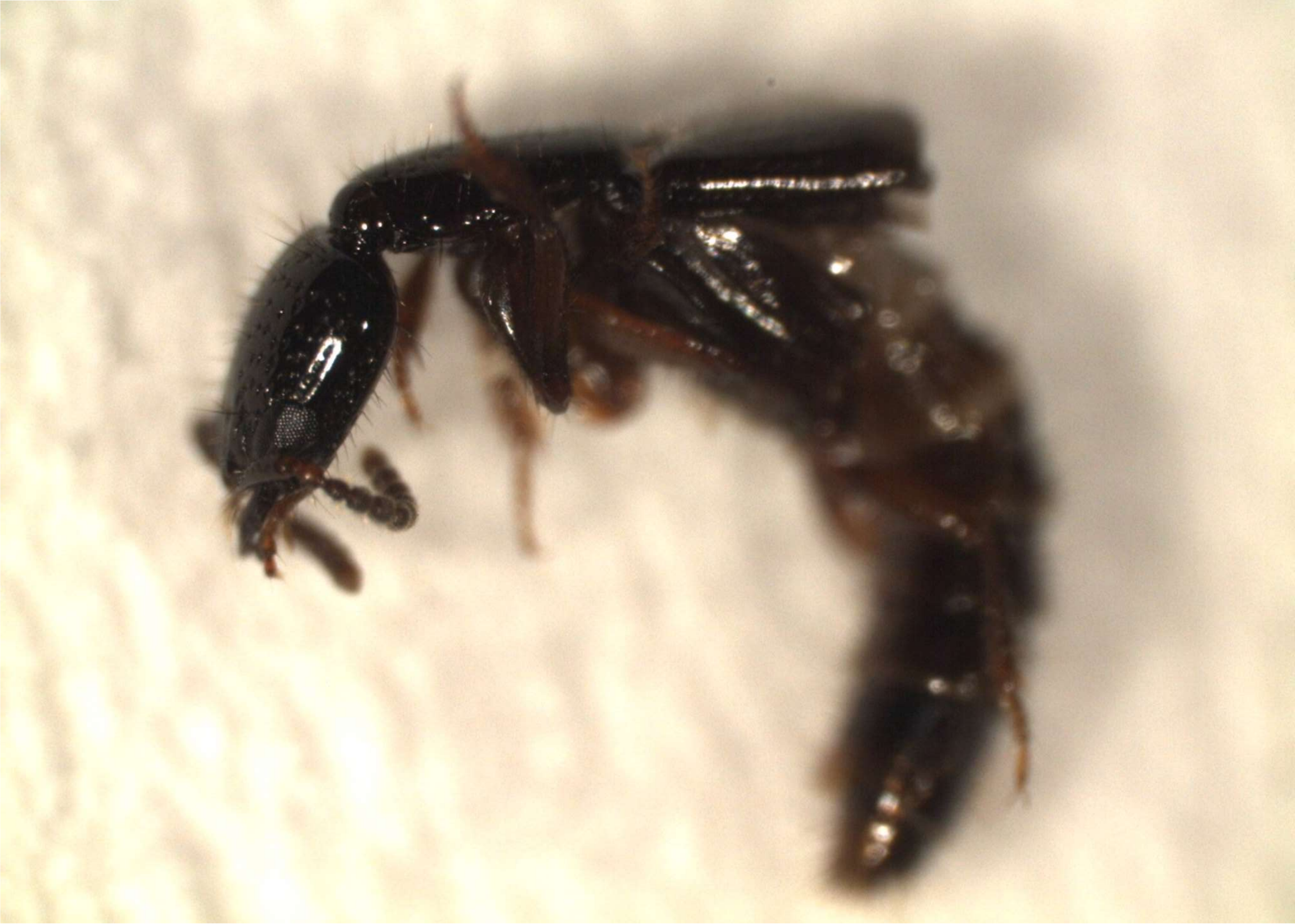

| Species FRE-6 |
| --- |
| 47-FRE-6 |

Non-myrmecophile – caught in forest litter away from army ant colonies

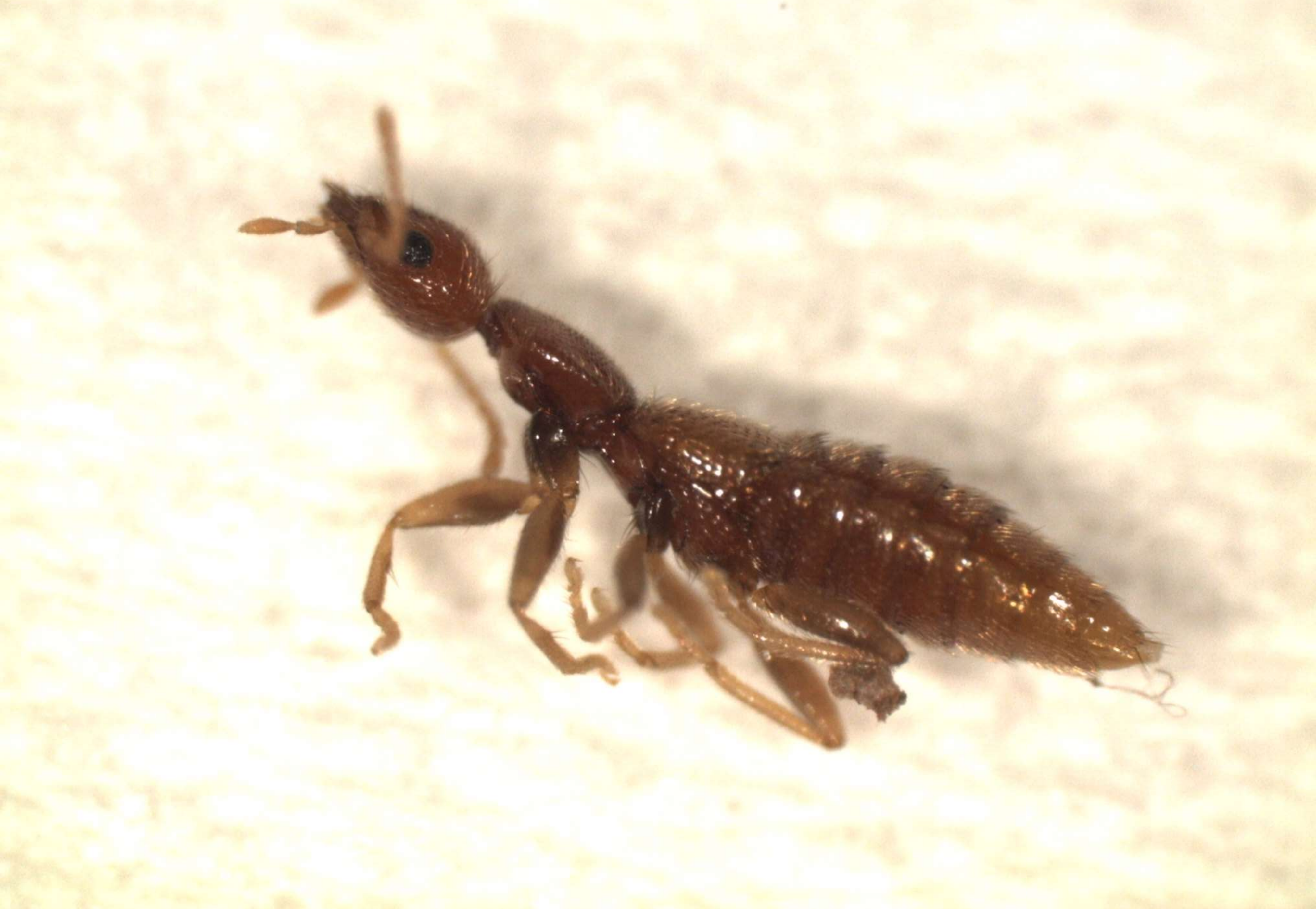
